# PTEN Deletion Robustly Boosts BRAF-Driven Intraspinal Sensory Axon Regeneration

**DOI:** 10.1101/2024.09.18.613685

**Authors:** Hyukmin Kim, Harun N. Noristani, Jinbin Zhai, Meredith Manire, Shuxin Li, Jian Zhong, Young-Jin Son

## Abstract

Primary sensory axons fail to regenerate into the spinal cord after dorsal root (DR) injury, resulting in persistent sensory deficits. This regenerative failure occurs at the dorsal root entry zone (DREZ), the CNS-PNS interface where injured sensory axons encounter both extrinsic inhibitory cues and a limited intrinsic growth state. Although several approaches have promoted partial DR regeneration across the DREZ, sustained long-distance regeneration, particularly of large-diameter myelinated axons, remains a major challenge. We previously showed that induced expression of constitutively active B-RAF (kaBRAF) increases the regenerative competence of injured adult DRG neurons. Here, we tested whether robust intraspinal regeneration after cervical DR injury could be achieved by selective kaBRAF expression alone or in combination with removal of myelin-associated inhibitors or deletion of neuron-intrinsic growth suppressors, PTEN or SOCS3. kaBRAF promoted reproducible but limited regeneration across the DREZ and did not produce significant functional recovery by two months. Additional deletion of Nogo, MAG, and OMgp produced only a modest improvement in kaBRAF-mediated regeneration. Deletion of PTEN or SOCS3, either alone or together, failed to promote meaningful growth across the DREZ. In contrast, PTEN deletion dramatically enhanced kaBRAF-mediated regeneration, enabling many axons to penetrate the DREZ and grow deep into the spinal cord, whereas SOCS3 deletion provided little additional benefit. These findings identify combined activation of BRAF-MEK-ERK and PI3K-Akt-mTOR signaling as a powerful strategy for stimulating robust intraspinal regeneration of injured DR axons.

## INTRODUCTION

After dorsal root (DR) injury, axons regenerate along the peripheral root but are largely prevented from re-entering the spinal cord at the dorsal root entry zone (DREZ), the CNS-PNS transition region 1. Failure of regenerating DR axons to reconnect with intraspinal targets can lead to persistent sensory loss, impaired sensorimotor coordination, and chronic pain despite deafferentation 2,3. Historically, regeneration failure at the DREZ has been attributed in large part to the inhibitory extracellular environment of the adult CNS 4-7. After injury, reactive astrocytes and the glial border are associated with increased chondroitin sulfate proteoglycans (CSPGs), whereas degenerating CNS myelin presents myelin-associated inhibitors such as Nogo, MAG, and OMgp. However, strategies that target these extrinsic inhibitors have produced only partial effects. For example, degradation of CSPGs with chondroitinase ABC induces plasticity and sprouting of intact cervical afferents rather than robust regeneration of injured axons 8, and genetic removal of major myelin-associated inhibitors has limited impact in several CNS injury models 39,54. Recent work on border-forming astrocytes after CNS injury further emphasizes that astrocytic boundaries are active, dynamic wound-repair structures rather than passive barriers alone 79.

A developmental decline in intrinsic growth capacity and limited trophic support after injury also contribute to regeneration failure at the DREZ 9-11. Exogenous neurotrophic factors can partially enhance DR regeneration by activating growth-associated signaling in adult DRG neurons. Intrathecal delivery of NGF, NT-3, or GDNF increases regeneration across the DREZ, but the response is largely restricted to axons that express the corresponding receptors 12,13.

Similarly, lentiviral expression of artemin or NGF in the dorsal horn promotes regeneration of small-diameter axons and improves nociceptive responses 14. Despite these advances, current strategies remain limited in their ability to promote robust regeneration across multiple sensory axon subtypes. In particular, there is still little compelling evidence for long-distance, functional regeneration of large-diameter proprioceptive afferents. In other CNS injury models, direct activation of intrinsic growth pathways by deleting PTEN or SOCS3, negative regulators of PI3K-Akt-mTOR and JAK-STAT3 signaling, respectively, promotes substantial axon regeneration 15-17. Whether these strategies can overcome the DREZ barrier after DR injury has remained unclear.

Neurotrophic factor activation of receptor tyrosine kinases engages RAF family serine/threonine kinases, including B-RAF, a potent activator of canonical MEK-ERK signaling 18-20. RAF signaling is required for neurotrophin-induced axon growth in embryonic DRG neurons 21,22, and kinase-active B-RAF promotes extensive growth of both TrkA-positive unmyelinated axons and parvalbumin-positive myelinated axons into the embryonic spinal cord 23. We previously showed that constitutively active B-RAF (kaBRAF; BRAFV600E) in adult DRG neurons promotes DR axon regeneration across the DREZ and into the dorsal horn within two weeks after DR crush 23,25,78. In the present study, we re-examined the effect of kaBRAF on sensory axon regeneration and tested whether its effects could be enhanced by deleting myelin-associated inhibitors or by genetically targeting PTEN and SOCS3. Using double and triple transgenic lines, we found that combined activation of BRAF-MEK-ERK and PI3K-Akt-mTOR signaling through kaBRAF expression and simultaneous PTEN deletion drives robust regeneration of large-diameter DR axons deep into the spinal cord within three weeks after injury.

## MATERIALS AND METHODS

### Animals

Male and female mice, 10-14 weeks of age and weighing 20-30 g, were used for all experiments. All animal care and procedures were conducted in accordance with the National Research Council’s Guide for the Care and Use of Laboratory Animals and were approved by the Institutional Animal Care and Use Committee at the Lewis Katz School of Medicine at Temple University, Philadelphia, PA, USA. Mouse transgenic lines were bred and maintained from lines generously provided by J. Zhong (Weill Cornell Medical College, NY), including LSL-kaBRAF:brn3a-CreERT2 (kaBRAF), PTENf/f:brn3a-CreERT2 (PTEN), SOCS3f/f:brn3a-CreERT2 (SOCS3), LSL-kaBRAF:SOCS3f/f:brn3a-CreERT2 (BRAF/SOCS3), LSL-kaBRAF:PTENf/f:brn3a-CreERT2 (BRAF/PTEN), PTENf/f:SOCS3f/f:brn3a-CreERT2 (PTEN/SOCS3), and LSL-kaBRAF:PTENf/f:SOCS3f/f:brn3a-CreERT2 (BRAF/PTEN/SOCS3).

The LSL-kaBRAF and brn3a-CreERT2 deleter mouse lines have been described previously 23-25. The LSL-kaBRAF:brn3a-CreERT2:Nogo-/-:MAG-/-:OMgp-/- (BRAF/tKO) line was generated by crossing the LSL-kaBRAF:brn3a-CreERT2 line with a Nogo/OMgp/MAG triple knockout line as described previously 26. Littermates were used as controls whenever possible.

Genotyping. Tail biopsies were obtained from 2- to 3-week-old mice. Samples were lysed and DNA was extracted using the REDExtract-N-AmpTM Tissue PCR kit (Sigma, #XNAT). DNA was amplified using the following primers for PCR: BRAF, 5’-GCC CAG GCT CTT TAT GAG AA-3’ (common forward), 5’-AGT CAA TCA TCC ACA GAG ACC T-3’ (reverse, mutant allele), 5’-GCT TGG CTG GAC GTA AAC TC-3’ (reverse, wildtype allele); TdTomato, 5’-AAG GGA GCT GCA GTG GAG TA-3’ (forward, wildtype allele), 5’-CCG AAA ATC TGT GGG AAG TC-3’ (reverse, wildtype), 5’-GGC ATT AAA GCA GCA TAT CC-3’ (reverse, mutant allele), 5’-CTG TTC CTG TAC GGC ATG G-3’ (forward, mutant allele); Brn3aCre, 5’-CGC GGA CTT TGC GAG TGT TTT GTG GA-3’ (forward), 5’-GTG AAA CAG CAT TGC TGT CAC TT-3’ (reverse); PTEN, 5’-CAA GCA CTC TGC GAA CTG AG-3’ (forward), 5’-AAG TTT TTG AAG GCA AGA TGC-3’ (reverse); SOCS3, 5’-CGG GCA GGG GAA GAG ACT GT-3’ (forward), 5’-GGA GCC AGC GTG GAT CTG C-3’ (reverse); Nogo, 5’-CAG TAG CTG CAG CAT CAT CG-3’ (common forward), 5’-CTC TCC AGC ACC TCC AAT TC-3’ (reverse, wildtype allele), 5’-AGA GGA ACT GCT TCC TTC AC-3’ (reverse, mutant allele); OMgp, 5’-GCA ATC AAC ATA AGA TGA CTT AAC-3’ (forward, wildtype allele), 5’-CAT TCT ATC ATA TAA AGG CTC CG-3’ (forward, mutant allele), 5’-ACA CAA CTT CTT CAC TCT CCC C-3’ (common reverse); and MAG, 5’-CTG CCG CTG TTT TGG ATA ATG-3’ (common forward), 5’-CGG AAA TAG TAT TTG CCT CCC-3’ (reverse, wildtype allele), 5’-ATG TGG AAT GTG TGC GAG GC-3’ (reverse, mutant allele).

4-Hydroxytamoxifen (4-HT) preparation. 4-HT stock (Sigma, H7904) was dissolved in 100% ethanol at 20 mg/ml. Sunflower seed oil (Sigma, S5007) was added to a 50 μl aliquot of 4-HT, and ethanol was evaporated by speed vacuum for 30 min. The sample was loaded into a 1 cc insulin syringe. Mice received 1 mg of 4-HT in sunflower seed oil by intraperitoneal injection once daily for 5 days. Control animals received sunflower seed oil alone. Mice were given a 2-day rest period before surgical procedures were performed.

Surgical procedures. One week after the first 4-HT injection, mice underwent dorsal root crush surgery as previously described 23,27. Briefly, mice were anesthetized by intraperitoneal injection of ketamine (80 mg/kg) and xylazine (12 mg/kg). Hair overlying the surgical site was removed with small animal clippers (Oster Professional Products). A 2-3 cm incision was made in the skin over the cervical region. The trapezius and paraspinal muscles, with associated fascia, were bluntly dissected and reflected using retractors to expose the dorsolateral aspects of the C2-T2 vertebrae. Using the spinous process of the T1 vertebra as a landmark, 2.5 mm curved rongeurs (Fine Science Tools) were used to make a series of right-sided hemilaminectomies and expose the C5-T1 spinal segments. Warm lactated Ringer’s solution was applied frequently to keep the exposed tissue moist. A 26-gauge subcutaneous needle (BD Biosciences) was used to make a small incision in the dura between each dorsal root. Lidocaine (2%) was added dropwise (∼2-3 drops) to the exposed spinal roots. Cervical roots were crushed for 10 s using fine forceps (Dumont #5) placed approximately 1-1.5 cm from the root insertion point. Muscles were sutured with sterile 5-0 sutures (Ethicon), and the midline incision was closed with wound clips (Fine Science Tools).

AAV vector injection. Recombinant adeno-associated virus (AAV) was used to label regenerating axons. At the time of dorsal root crush or at least two weeks before sacrifice, the C5, C6, and/or C7 cervical DRGs were exposed. Animals were placed in a stereotaxic holder with spinal cord clamps (STS-A, Narishige Group, Japan) to stabilize the spinal cord and DRGs during injection. AAV2-GFP (Vector Biolabs, Cat. #7072, 1 x 10^13 GC/ml), which expresses eGFP under the control of a CAG promoter, was injected into the cervical DRGs (800-1000 nl total volume; ∼100 nl/min) using a glass micropipette and nano-injector (World Precision Instruments, Sarasota, FL). The micropipette was left in place for 3-5 min after each injection to ensure diffusion into the DRG. After injection, a piece of biosynthetic membrane (Biobrane, UDL Laboratories, Sugarland, TX) was placed over the exposed spinal cord. Muscles were closed with sterile 5-0 silk sutures (Ethicon), and the midline skin incision was closed with wound clips 28. Animals were placed on a heating pad during recovery from anesthesia and received buprenorphine (0.05 mg/kg) intramuscularly for postoperative analgesia.

Behavioral analysis. All behavioral experiments were performed and scored by an observer blinded to experimental group. Testing was performed before 4-HT injection or at least one day before DR crush and then at 7-day intervals after DR injury for up to two months. All tests were performed on the same day. Recovery of sensory and sensorimotor function was assessed using the horizontal ladder test, beam walking test, von Frey test of mechanical sensitivity, forepaw sticker test, paw-withdrawal latency to noxious thermal stimulation, and grip strength.

Horizontal ladder test. The horizontal ladder test was used to assess proprioceptive recovery as previously described ^29,30^. Mice were placed on a horizontal ladder with 11 cm wide rungs spaced 2 cm apart. The ladder was elevated about 1 meter off the ground and tilted upward slightly to encourage locomotion. The side walls of the ladder were 7 cm high. During the test, mice walked from a neutral starting box towards a darkened box that contained bedding material taken from their respective home cage. The mice were required to walk at least 66 cm along the ladder per trial, three trials total. The total number of steps and the total number of foot missteps or slips from both the ipsilateral and contralateral paws were counted. All trials were video recorded and analyzed at a later time point to ensure accurate counts of steps and slips.

Narrowing beam walking test. Mice traversed a 1 m long by 10 cm high wooden beam that progressively narrowed from 3 cm at the starting point to 5 mm at the end. A cereal treat was placed at the end of the beam to encourage mice to walk the full length. Ipsilateral forepaw performance was scored as follows: 0, no attempt to use the limb; 1, movement without weight bearing; 2, attempted weight bearing with frequent missteps and walking on the dorsal surface of the paw; 3, weight bearing with fewer missteps or misplacements; 4, normal weight bearing with very few mistakes; and 5, normal walking without deficits. This scoring system was adapted from a previously published scale 31. Scores were averaged over three trials. The distance to first slip of the right forepaw off the beam was also scored and is presented as units before first slip, with one unit equal to 5 cm.

Von Frey filament test of mechanical sensitivity. Mice were placed in an elevated transparent plastic box with a mesh metal floor (8 x 12 x 10 cm) and allowed to acclimate for at least 1 h. Mechanical sensitivity was measured using von Frey filaments with bending forces of 0.02 g, 0.04 g, 0.4 g, 1 g, 4 g, and 8 g (Touch Test Sensory Evaluator, Stoelting, Wood Dale, IL). The up-and-down method was used to measure the 50% paw withdrawal threshold 32,33. Beginning with the 1 g filament, the sequence of responses to increasing or decreasing monofilaments was recorded. Filaments were applied perpendicularly to the mid-plantar region of the ipsilateral or contralateral forepaw in one steady motion with enough pressure to bend the filament without lifting the paw from the mesh floor. A positive response was defined as a rapid lifting of the forepaw while the filament was in contact with the plantar surface (<2 s). If no response was observed, the next higher filament was applied; if a positive response was observed, the next lower filament was applied. Consecutive stimuli were separated by 30 s to avoid adaptation. Each animal was assigned a tabular value (k) based on the response pattern 32. The 50% response threshold was calculated using the formula: 50% g threshold = (10[Xf+kδ])/10,000, where Xf is the value of the last von Frey hair used, k is the tabular value, and δ is the mean difference between stimuli in log units (0.520). Threshold values were calculated for both ipsilateral and contralateral forepaws.

Hot water test. Nociceptive recovery was assessed by measuring response latency to noxious thermal stimulation, a 50 °C water bath ^34^. Both the contralateral and ipsilateral forepaws were tested. Mice were lightly restrained in a cloth harness to allow for isolation of the forelimb and to prevent unintentional damage to the injured cervical region that would result from scruffing. The protruding forepaw was then dipped into the hot water bath up to the wrist area. The latency to response, or time to removal of the forepaw from the water, was measured to the nearest 0.01 seconds. A maximum cut-off time of 20 seconds was used to prevent unnecessary damage to the paw. Three trials were performed per paw, separated by >30 seconds, and averaged. Careful attention was paid to ensure that the whiskers and tail did not touch the hot water during the trials.

Immunohistochemistry. Three or ten weeks after injury, mice were anesthetized with a lethal dose of 10% Euthasol (Virbac, USA) in 0.9% saline. Mice were transcardially perfused with a 4% (vol/vol) solution of paraformaldehyde in 0.01 M PBS, pH 7.4. The cervical spinal cord and dorsal root ganglia were dissected out and post-fixed in 4% paraformaldehyde for 2 hours. The tissue was cryoprotected in 30% sucrose (wt/vol) in 0.01 M PBS at 4 °C for 24 hours. Tissue was placed in a cryomold filled with tissue embedding matrix (M-1 Embedding Matrix, Thermo Fisher) and quickly immersed in a beaker of 2-methylbutane cooled with liquid nitrogen. The tissue was allowed to freeze completely prior to removal. The frozen tissue was sectioned at 20 μm and collected on slides. For immunostaining, sections were rehydrated in 0.01 M PBS, incubated in 0.1 M glycine/2% bovine serum albumin (BSA)/PBS, and then blocked with 0.2% Triton X-100 in 2% BSA/PBS. Primary antibodies for immunostaining were used at the following dilutions: rabbit anti-NF200 (1:500, Sigma, #N4142), IB4-biotin conjugate (1:200, Sigma, L2140), rabbit anti-CGRP (1:1000, Peninsula Labs, CA, #T-4032), chicken anti-GFP (1:1000, Avés Labs Inc., #1020), rabbit anti-GFAP (1:500, Dako, Z0334), and rabbit anti-PTEN (1:200, Cell Signaling, #9559). Sections were incubated in primary antibodies overnight at 4 °C. Sections were washed in 2% BSA/PBS and then incubated for 1-2 hours at room temperature in secondary antibody. Secondary antibodies used were Alexa Fluor 647 goat anti-rabbit IgG (1:400, Molecular Probes, A-21244), Fluorescein (FITC)-conjugated goat anti-rabbit IgG (1:400, Chemicon International, AP307F), rhodamine (TRITC) streptavidin (1:400, Jackson ImmunoResearch Labs Inc., 016-020-084), or Alexa Fluor 488 donkey anti-chicken IgG (1:400, Jackson ImmunoResearch Labs Inc., 703-545-155). DAPI nucleic acid stain (1:1000, Invitrogen, D-1306) for neuronal or non-neuronal cell nuclei or Nissl substance stain (1:200, Molecular Probes, N-21479) for neuronal cells were used to counterstain prior to final wash steps in 0.01 M PBS. Stained sections were mounted with media for fluorescence (Vectashield, Vector Laboratories Inc., CA) and a glass coverslip.

Western blots. Mice were deeply anesthetized as described above and decapitated. The left and right C1-L5 DRGs and roots were quickly collected and washed with Hank’s buffered salt solution (HBSS) or 0.01 M PBS. Excess buffer was removed, and tissue was flash frozen in liquid nitrogen and stored at −80 °C until processing. Samples were lysed on ice in RIPA buffer (50 mM Tris-HCl, pH 8.0; 150 mM sodium chloride; 0.5% Triton X-100; 0.5% sodium deoxycholate; 0.1% sodium dodecyl sulfate [SDS]) containing protease and phosphatase inhibitors. Protein concentration was determined using a bicinchoninic acid (BCA) protein assay kit (Thermo Fisher, #23227). Lysate was mixed with 4X NuPAGE LDS sample buffer and 10X sample reducing agent (Thermo Fisher) to obtain a 1X solution containing 25-50 μg protein.

Lysates were resolved on 7.5, 8, 10, or 12% SDS-PAGE gels and transferred to nitrocellulose membranes. Membranes were blocked in 5% milk/TBST for 1 h at room temperature and incubated with primary antibodies in 1% milk/TBST overnight at 4 °C. Primary antibodies were mouse anti-BRAF V600E (1:1000, Spring Bioscience), rabbit anti-BRAF (1:1000, Cell Signaling #9433), rabbit anti-PTEN (1:1000, Cell Signaling #9559), rabbit anti-pS6 (1:1000, Cell Signaling #2215), rabbit anti-survivin (1:1000, Cell Signaling #2808), and mouse anti-β-actin (1:5000, Sigma, A5441). Blots were washed with TBST and incubated with HRP-linked donkey anti-mouse IgG (1:5000, GE Healthcare) or HRP-linked donkey anti-rabbit IgG (1:5000, Jackson ImmunoResearch) for 1 h at room temperature. Blots were developed using ECL (GE Healthcare).

Microscopy and image acquisition. An Olympus BX53 fluorescence microscope with an X-Cite 120Q light source (Lumen Dynamics) was used to visualize immunostained tissue sections. The microscope was equipped with an ORCA-R2 digital CCD camera (Hamamatsu) controlled by MetaMorph Image Analysis Software (Molecular Devices). Z-stack images were acquired using a Zeiss Axio Imager upright fluorescence microscope with AxioVision software or a Leica SP8 confocal microscope with a 20X objective and Leica software. Images were processed using Imaris (Bitplane) and Photoshop (Adobe).

Axon density and axon number index. Axon regeneration was quantified by measuring both the axon number index and axon density within the spinal cord. For the axon number index, images from GFP-labeled sections were acquired, converted to 8-bit in ImageJ (NIH), and filtered using the FeatureJ Hessian plugin to obtain smallest-eigenvalue images 35. Using the astrocyte border, demarcated by GFAP immunostaining, as a marker of the DREZ, lines were drawn at 100 μm before the DREZ (within PNS territory), at the DREZ (GFAP outer boundary), and at 100 μm intervals into CNS territory. The number of axons at each distance was counted using the ImageJ line profile tool. If axons could not be resolved using ImageJ, they were counted manually.

Initial counts were normalized to the number of labeled fibers in the dorsal root, and data are presented as the ratio of labeled axons at each distance past the DREZ to labeled axons in the dorsal root. For axon density, images were converted to 8-bit in ImageJ and filtered using the FeatureJ Hessian plugin to obtain median-eigenvalue images. Optical density of GFP-labeled axons within spinal cord gray matter was measured and normalized to optical density within the DR, using the rectangular selection tool to draw a consistently sized box beginning 100 μm into the DR from the GFAP border. Data are presented as relative pixel intensity. At least five sections within the injury area were selected at random from at least three animals for both measures of axon regeneration.

Statistical analysis. GraphPad Prism software was used for all statistical analyses. Data from AAV2-GFP labeling were analyzed using an unpaired Student’s t-test assuming Gaussian distribution. Control and experimental group data from behavioral assessments were compared using two-way ANOVA followed by Tukey’s post hoc test. Axon density and axon number index were analyzed by one-way or two-way ANOVA with Tukey’s or Sidak’s post hoc tests, as indicated in the figure legends. All data are presented as mean ± SEM. Sample sizes are reported in the figure legends. Results were considered statistically significant when P < 0.05.

## RESULTS

### AAV2-GFP selectively labels large-diameter NF+ dorsal root axons

Viral tracers can overcome several limitations of conventional axon tracing, including nonspecific labeling and difficulty distinguishing spared axons from regenerating or sprouting axons 36. Recombinant adeno-associated virus (AAV) provides efficient neuronal transduction with low immunogenicity and minimal host cytotoxicity 37,38. We therefore used AAV serotype 2 to label sensory axons with enhanced green fluorescent protein (eGFP). We first characterized transduction efficiency and neuronal tropism after AAV2-eGFP microinjection into cervical DRGs of uninjured adult wildtype mice. GFP expression was detected in most DRG neuronal cell bodies, indicating efficient transduction of sensory neurons (Figure 1A). Quantification showed that 83 ± 11% (497/581) of the total sensory neuron population was transduced by AAV2-eGFP (Figure 1B). GFP expression was not detected in non-neuronal or glial cell populations, and all GFP+ cells co-labeled with Nissl.

**Figure 1.**
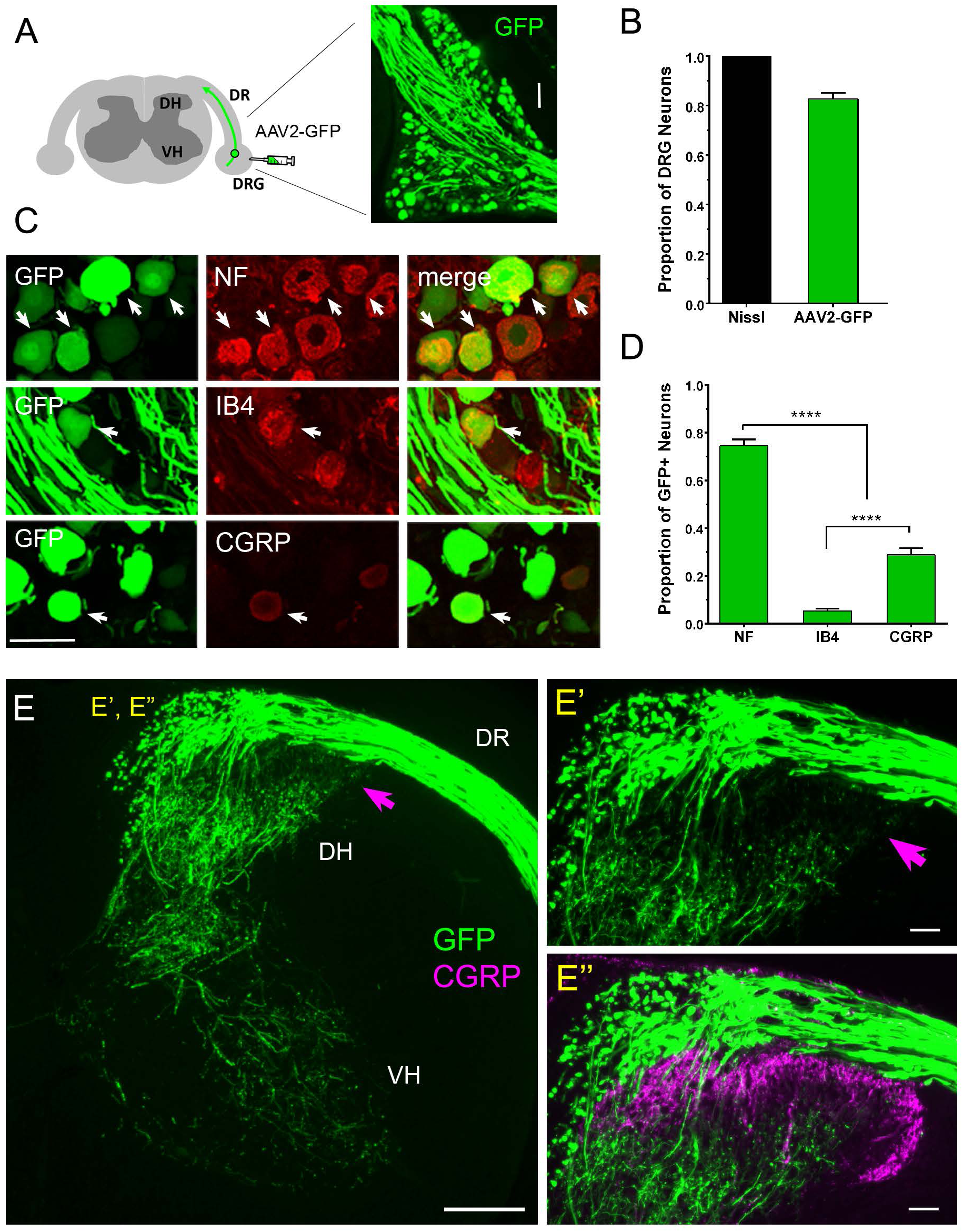
AAV2-GFP selectively labels large-diameter axons of the DRG. (A) Two weeks after intraganglionic injection of AAV2-GFP, GFP is visible in many sensory neurons of the adult wildtype DRG. Scale bar: 100 μm. (B) Quantification of neuronal transduction efficiency shows GFP expression in the majority of Nissl+ sensory neurons. Nissl was used to label the total neuronal population. (C) Immunostaining shows GFP co-labeling with NF+, IB4+, and CGRP+ sensory neurons. Arrows indicate co-labeled cells. Scale bar: 50 μm. (D) Analysis of AAV2-GFP sensory subtype tropism shows that AAV2-GFP+ neurons are predominantly NF+. **** P < 0.0001, one-way ANOVA. n = 3 mice, 5-8 sections per mouse, separated by at least 100 μm. Values represent mean ± SEM. (E) GFP+ sensory fibers in the DR project to the dorsal and ventral horns in intact animals. GFP+ afferents are sparse in the most superficial laminae of the DH (E’), corresponding to areas containing CGRP+ small-diameter axons (E’’). Pink arrows indicate areas of minimal GFP labeling. Scale bars: E = 200 μm; E’ and E’’ = 50 μm. DR = dorsal root; DH = dorsal horn; VH = ventral horn.

DRGs contain heterogeneous sensory neurons that can be broadly grouped into large-diameter neurons, small-diameter peptidergic neurons, and small-diameter non-peptidergic neurons. We next asked whether AAV2-eGFP transduced all DRG neuron subtypes or showed subtype preference. Immunohistochemical analysis showed GFP co-localization with markers of both large- and small-diameter sensory neurons (Figure 1C). Quantification revealed that 74 ± 13% (370/497) of GFP+ neurons co-labeled with the large-diameter marker NF, 6 ± 4% (65/1181) with the small-diameter non-peptidergic marker IB4, and 29 ± 12% (328/1181) with the small-diameter peptidergic marker CGRP (Figure 1D). Significantly more GFP+ neurons co-labeled with NF than with either IB4 or CGRP. Thus, AAV2 transduces multiple sensory neuron subtypes but displays a clear preference for large-diameter neurons, including proprioceptors and mechanoreceptors.

Consistent with this cell-body tropism, robust GFP expression was observed in both peripheral and central processes of transduced sensory neurons two weeks after injection (Figure 1A, E). GFP-labeled fibers were visible along the DR and throughout the spinal cord in patterns consistent with known afferent trajectories (Figure 1E). However, fewer GFP-labeled fibers were observed in laminae I and II of the superficial dorsal horn, where small-diameter nociceptive axons, including CGRP+ fibers, normally terminate (Figure 1E’-E’’). These findings confirm that intraganglionic AAV2-eGFP injection is an effective approach for labeling large-caliber DR axons. In the experiments below, AAV2-eGFP labeling was used to assess regeneration of large-diameter DR axons into the spinal cord.

### kaBRAF promotes penetration of axons across the DREZ

We previously showed that conditional expression of kinase-active B-RAF (kaBRAF) enables many adult DR axons to regenerate across the DREZ and into the dorsal horn within two weeks after injury 23. Subsequent observations from our laboratory indicated that inflammatory lesions at or near the DREZ can increase apparent intraspinal growth independently of transgene activation, potentially confounding anatomical interpretation. To reassess kaBRAF-mediated regeneration under more stringent conditions, we excluded spinal cords with inflammatory disruption of the GFAP+ astrocytic boundary near the ipsilateral DREZ. The experimental design is summarized in Figure 2A. Vehicle or 4-HT was administered for 5 days, multi-root crush was performed on day 8, and tissue was collected 21 days later. Before tissue processing, spinal cords were examined in wholemount to confirm complete crush injury and absence of spared axons; spinal cords with incomplete crushes were excluded.

**Figure 2.**
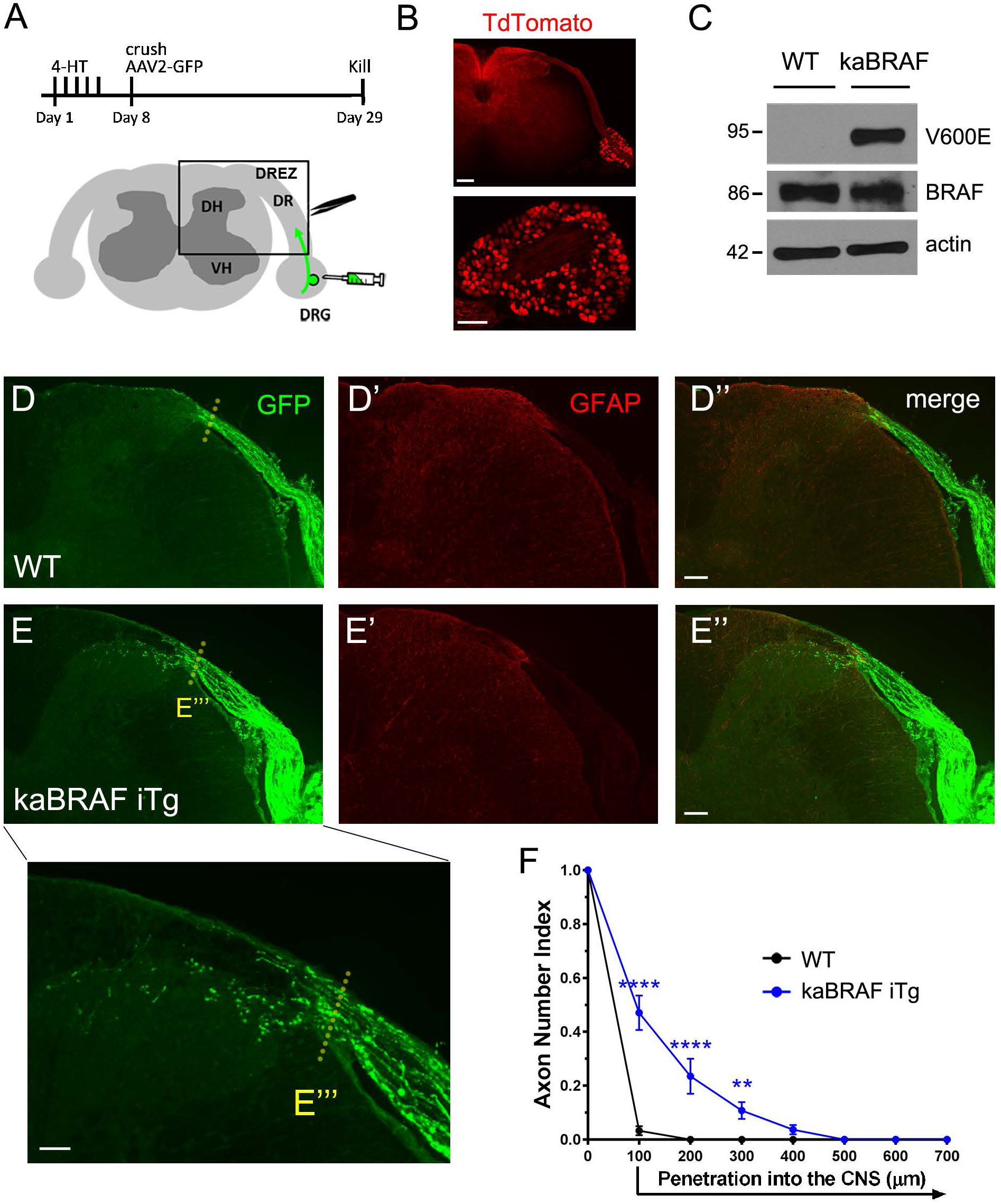
kaBRAF enables penetration of regenerating axons across the DREZ. (A) Schematic diagram of the experimental procedure for 3-week analysis of transgenic lines. Boxed area indicates the region of interest in the spinal cord. (B) tdTomato expression indicates high transgene activation within sensory neurons and afferent projections in the dorsal horn and dorsal columns. Scale bars: upper panel = 200 μm; lower panel = 100 μm. (C) Western blot analysis confirms expression of BRAFV600E in kaBRAF iTg but not WT DRG lysates. Total B-RAF protein levels remain unchanged. Molecular mass is indicated in kilodaltons. Cross sections from WT (D-D’’) and kaBRAF iTg (E-E’’’) mice show GFP-labeled axon regeneration 3 weeks after DR crush injury. WT axons regenerate along the DR but do not enter the spinal cord. (E’’’, inset) kaBRAF expression promotes axon penetration across the DREZ and into superficial spinal cord laminae. GFAP staining denotes the astrocytic boundary of the CNS. Dotted lines indicate the approximate DREZ. (F) Quantification of axon growth shows significantly greater DREZ penetration in kaBRAF iTg than in WT controls. Scale bars: D-E’’ = 100 μm; E’’’ = 50 μm. n = 4-5 mice per group, at least 5-6 sections per mouse. **** P < 0.0001, ** P < 0.01, two-way ANOVA with Sidak’s multiple comparisons test. Values represent mean ± SEM.

We used the LSL-kaBRAF:tdTomato:brn3aCreERT2 gain-of-function mouse line to conditionally express kaBRAF in adult DRG sensory neurons as previously described 23. After 4-HT administration, tdTomato fluorescence was detected in nearly all (>90%) sensory neurons of the DRG, indicating efficient Cre recombination and transgene activation (Figure 2B).

Western blot analysis confirmed expression of BRAFV600E in DRG lysates (Figure 2C). Total B-RAF protein levels were not detectably altered between wildtype and kaBRAF samples, consistent with expression of kaBRAF from the endogenous B-RAF locus.

We next determined whether kaBRAF expression enabled injured DR axons to cross the DREZ and enter the spinal cord. In wildtype animals, injured DR axons regenerated along the DR but failed to cross the GFAP+ astrocyte boundary into the CNS (Figure 2D-D’’). In contrast, kaBRAF expression modestly enhanced regeneration across the DREZ, allowing a subset of axons to enter the superficial laminae of the dorsal horn by 3 weeks after DR injury (Figure 2E-E’’’). This increase was consistently observed across injured roots. Quantification showed significant differences between wildtype and kaBRAF groups at 100 μm (P < 0.0001) and 200 μm (P = 0.0007) central to the GFAP border (Figure 2F). Nevertheless, most axons still failed to penetrate beyond the DREZ, and only rare axons extended deeply into CNS territory. Thus, kaBRAF enhances the intrinsic growth capacity of injured sensory neurons but is not sufficient to drive robust long-distance intraspinal regeneration.

### kaBRAF sustains axon growth over two months but fails to elicit robust intraspinal regeneration or functional recovery

To determine whether kaBRAF expression supports continued axon growth over a longer period, we assessed anatomical regeneration and behavioral recovery two months after DR injury. A C5-T1 DR crush was performed to remove forelimb afferent input, and AAV2-eGFP was injected into C5-C7 DRGs two weeks before sacrifice. In control animals, few axons were found within the DREZ territory, and most remained stalled at the astrocyte border (Figure 3A-A’). Compared with the 3-week time point, 2-month kaBRAF expression allowed a similar number of axons to cross the GFAP border, but more of these axons extended deeper into the DREZ and superficial dorsal horn (Figure 3B-C). Regenerating axons remained confined to the gray matter. Axon number index analysis showed a significant difference between 3-week and 2-month kaBRAF groups at 200 μm, but not at 100 μm, central to the GFAP border (Figure 3C).

**Figure 3.**
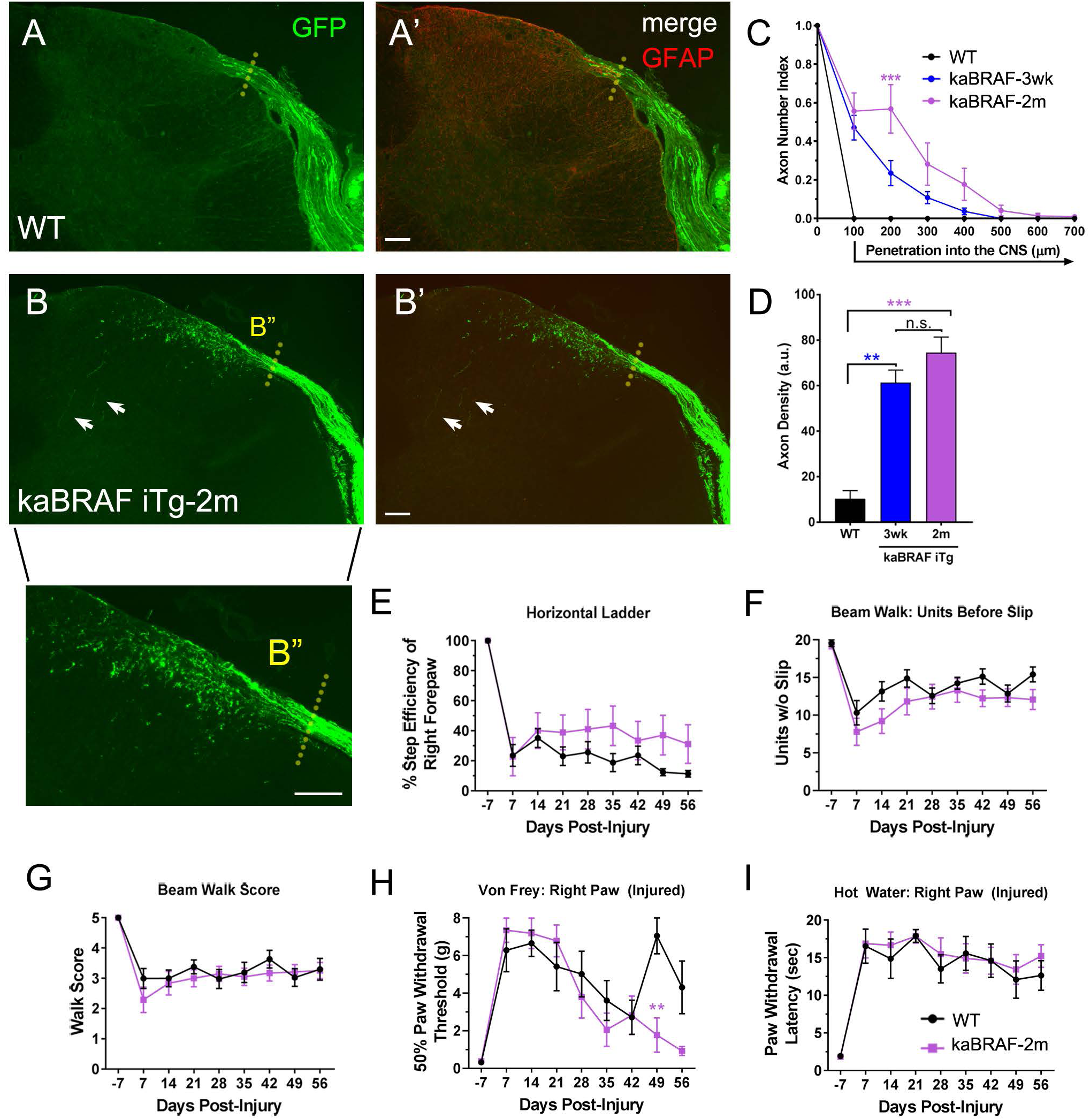
kaBRAF sustains axon regeneration but does not lead to functional recovery by 2 months after injury. Cross sections show GFP-labeled sensory axon regeneration 10 weeks after DR crush injury in WT and kaBRAF iTg mice. (A-A’) Regenerating axons of WT mice remain within the DREZ territory even at 10 weeks. (B-B’) kaBRAF expression continues to promote axon growth into the superficial spinal cord, with greater penetration than observed at 3 weeks (Fig. 2E-E’’’). A few axons are observed in deeper laminae (arrows). (B’’) Higher-magnification view of regenerating axons. Scale bar: 100 μm. GFAP staining denotes the astrocytic boundary of the CNS. Dotted lines indicate the approximate DREZ. (C) Quantification of axon growth shows a significant difference between kaBRAF iTg 2-month and 3-week groups at 200 μm from the GFAP border. Compared with WT, kaBRAF iTg 2-month mice also show significant axon growth at 200 μm (****) and 300 μm (**). (D) Axon density analysis shows a significant difference between WT and kaBRAF iTg groups. n = 3-6 mice per group, at least 5-6 sections per mouse. **** P < 0.0001, *** P < 0.001, ** P < 0.01, n.s. = not significant, one-way or two-way ANOVA with Tukey’s multiple comparisons test. kaBRAF does not significantly enhance recovery of proprioceptive function (E-G), response to mechanical stimuli (H), or response to noxious thermal stimuli (I) compared with WT controls. No significant difference was observed between groups at any time point in horizontal ladder walk, beam walk, or hot water testing. A minor but significant difference was observed at day 49 in the von Frey filament test. n = 8 mice per group, ** P = 0.002, two-way ANOVA with Sidak’s multiple comparisons test. Values represent mean ± SEM.

Axon density within the spinal cord did not differ between 3-week and 2-month kaBRAF groups (Figure 3D). In wholemount and transverse preparations, GFP-labeled regenerating axons were observed in gray matter at cervical levels C4 and C3 but were absent from C8 (not shown), suggesting that kaBRAF-driven growth may retain some rostrocaudal specificity.

We then asked whether this sustained anatomical regeneration was sufficient to improve proprioceptive, tactile, or nociceptive function. Behavioral recovery was assessed using horizontal ladder walk, beam walk, von Frey filament testing, and hot water withdrawal.

Proprioceptive recovery is particularly difficult after DR injury because regenerating axons must travel long distances to reconnect with second-order neurons. Prior to injury, control and kaBRAF mice showed 100% efficiency of right and left forepaw placement on the horizontal ladder (Figure 3E). Seven days after C5-T1 DR crush, right forepaw placement efficiency dropped markedly in both control (23.5 ± 7.3%) and kaBRAF (22.7 ± 12.8%) groups relative to baseline. Two-way ANOVA revealed no significant difference between groups over time.

Similarly, beam walk analysis showed no significant difference between groups in either distance traveled before forepaw slip (Figure 3F) or injured forepaw placement score (Figure 3G). Mechanical sensitivity, measured by von Frey testing, did not differ significantly between control and kaBRAF groups at any time point except day 49 (Figure 3H).

Thermal nociception was assessed by measuring paw withdrawal latency from a 50 °C water bath. Both control and kaBRAF mice showed loss of nociceptive function in the injured right forepaw 7 days after C5-T1 DR crush, reflected by increased withdrawal latency.

Nociceptive responses in the uninjured forepaw remained stable throughout the testing period. Neither group showed pronounced restoration of thermal nociception by two months after injury, and two-way ANOVA revealed no significant difference between groups (Figure 3I). These findings indicate that although kaBRAF sustains limited intraspinal growth over time, it is not sufficient to restore sensory function in this injury model.

### Additional deletion of myelin-associated inhibitors modestly enhances kaBRAF-elicited intraspinal regeneration

Because adult CNS myelin contains inhibitory molecules that can restrict axon growth 39, we next asked whether removal of Nogo, MAG, and OMgp could enhance kaBRAF-mediated regeneration at 3 weeks after DR injury. We crossed the LSL-kaBRAF:tdTomato:brn3aCreERT2 line with a Nogo-/-:MAG-/-:OMgp-/- triple knockout line to generate kaBRAF/tKO mutants 26,39. Triple deletion of Nogo, MAG, and OMgp alone did not enhance axon regeneration across the DREZ compared with wildtype controls (Figure 4A-A’, C, D). In contrast, combined deletion of the myelin-associated inhibitors with kaBRAF expression increased regeneration across the DREZ compared with tKO or kaBRAF alone (Figure 4B-D). Quantification showed significant differences in axon number index between tKO and kaBRAF/tKO groups at 100 and 200 μm central to the GFAP border (Figure 4C). Axon density also differed significantly between tKO and kaBRAF/tKO groups (Figure 4D). However, compared with kaBRAF alone, kaBRAF/tKO produced only a modest improvement, with a significant difference in axon number index at 100 μm but no significant difference in overall axon density. Thus, removing major myelin-associated inhibitors modestly improves the response to kaBRAF but is not sufficient to drive robust intraspinal regeneration.

**Figure 4.**
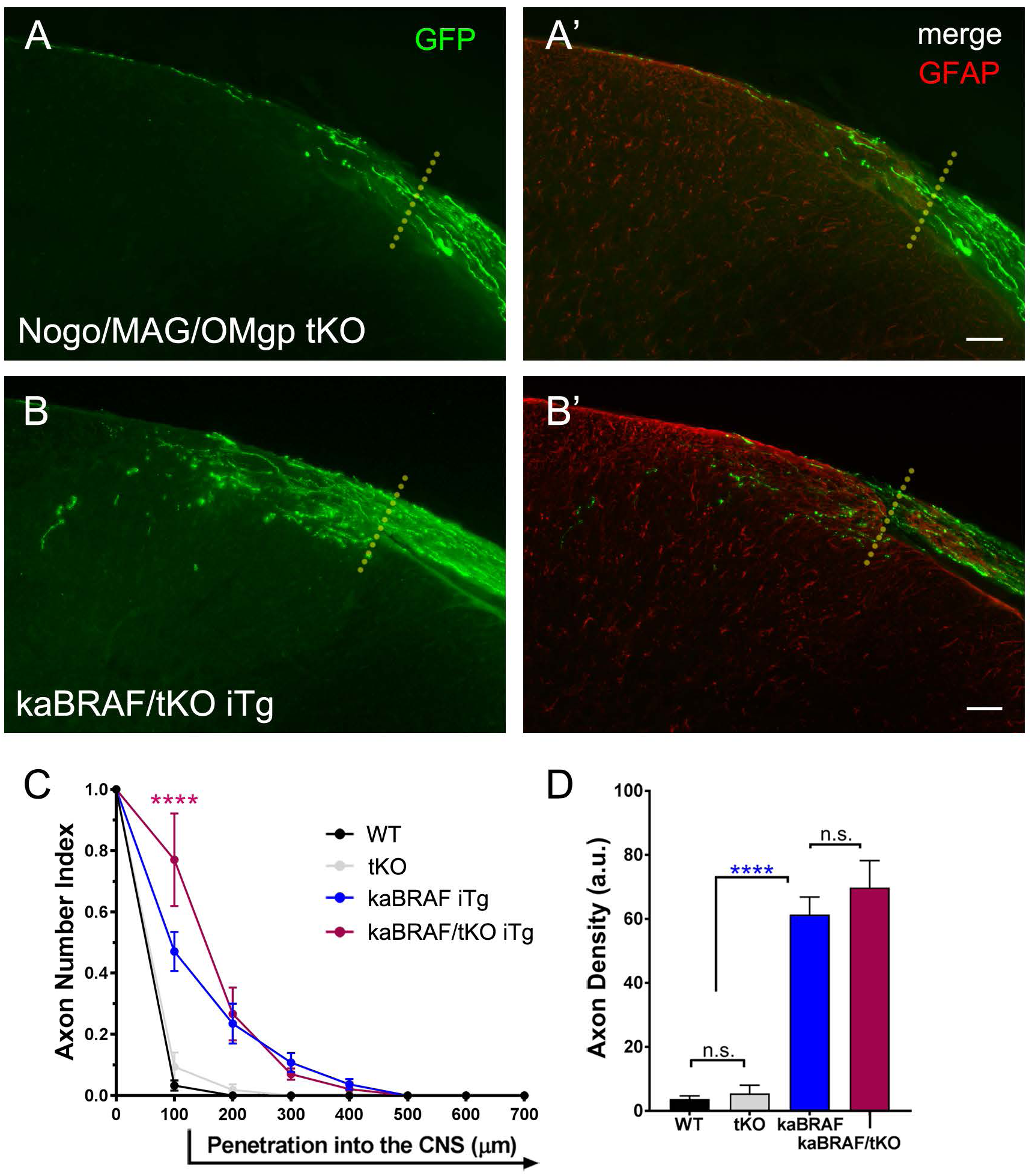
Additional deletion of myelin-associated inhibitors minimally improves kaBRAF-mediated axon regeneration at 3 weeks after injury. Cross sections show GFP-labeled regenerating sensory axons in tKO (A-A’) and kaBRAF/tKO iTg (B-B’) mice 3 weeks after DR crush injury. GFAP staining denotes the astrocytic boundary of the CNS. Dotted lines indicate the approximate DREZ. Scale bar: 50 μm. (C) Quantification of axon growth shows significantly improved regeneration in the kaBRAF/tKO group compared with the kaBRAF group at 100 μm from the GFAP border. The kaBRAF/tKO group also shows significant growth compared with tKO at 100 μm (****) and 200 μm (***). No significant difference is observed between tKO and WT groups. (D) Axon density shows no significant difference between kaBRAF and kaBRAF/tKO groups but shows significant differences between kaBRAF or kaBRAF/tKO groups and tKO or WT groups. n = 3-5 mice per group, at least 5-6 sections per mouse. **** P < 0.0001, *** P < 0.001, n.s. = not significant, one-way or two-way ANOVA with Tukey’s multiple comparisons test. Values represent mean ± SEM.

### PTEN deletion does not enable axons to penetrate the DREZ

PTEN is a major negative regulator of the PI3K-Akt-mTOR growth-associated signaling pathway 40. Conditional PTEN deletion promotes optic nerve and corticospinal tract regeneration through mTOR-dependent and mTOR-independent mechanisms 17,41, but its effect in the DR injury model has not been directly tested. We therefore used a brn3aCreERT2:tdTomato:PTENf/f transgenic line to conditionally delete PTEN from adult DRG sensory neurons. Because PTEN expression in adult sensory neurons has been less extensively characterized, we first confirmed its baseline expression in wildtype DRGs.

Immunohistochemistry showed widespread PTEN expression in DRG neurons and particularly strong expression in small-diameter neurons, consistent with a previous report 42 (Figure 5A). PTEN levels were similar in intact and injured wildtype DRGs two weeks after injury (Figure 5A). Cre-mediated recombination substantially reduced PTEN expression, as confirmed by immunohistochemistry and Western blotting (Figure 5A, B). PTEN deletion also increased phospho-S6 (pS6), indicating activation of PI3K-Akt-mTOR signaling (Figure 5B).

**Figure 5.**
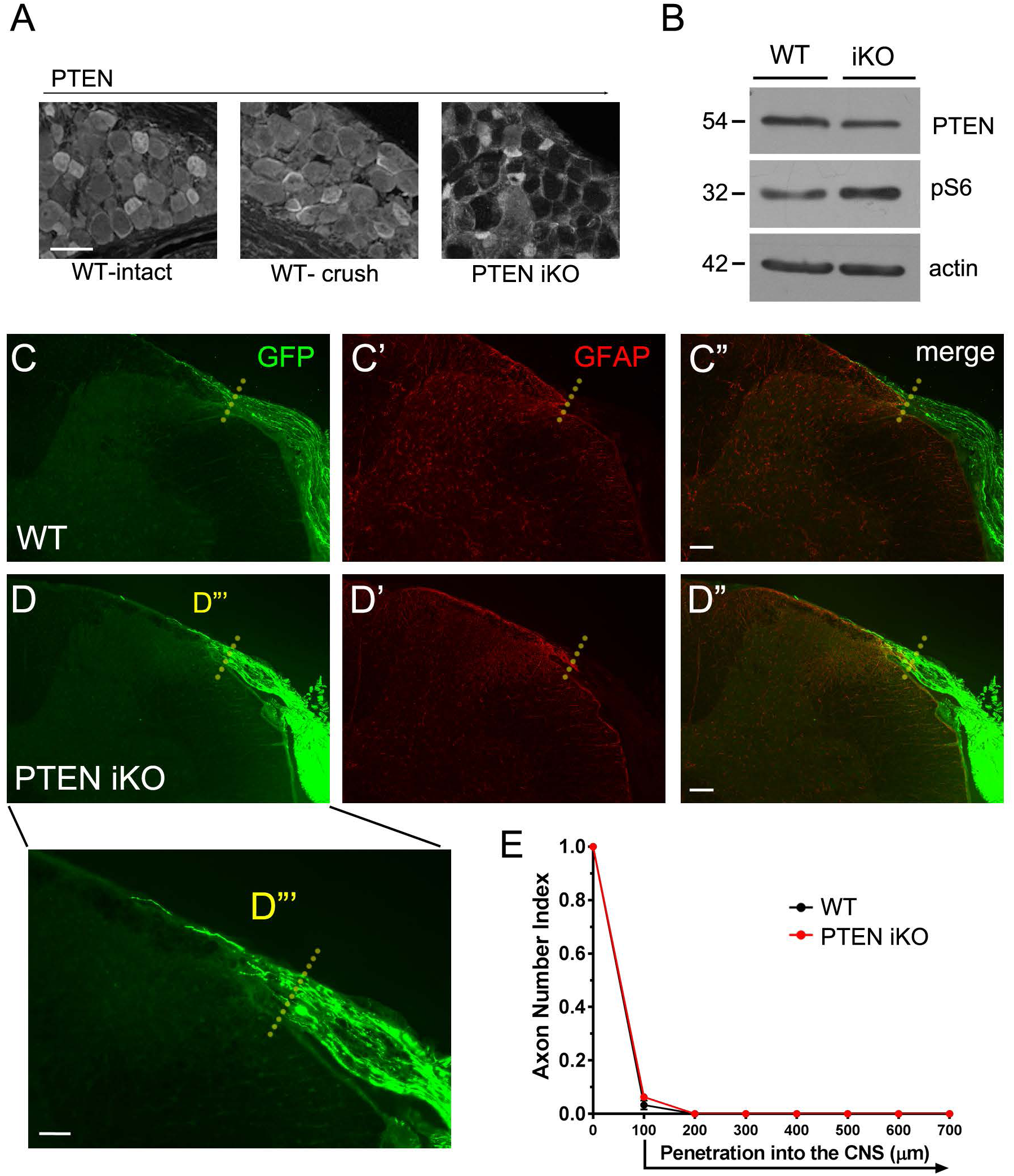
PTEN deletion does not promote axon regeneration across the DREZ. (A) Immunostaining of adult DRG for PTEN shows similar levels in intact and injured WT mice (3 weeks post-crush), with particularly high levels in small-diameter neurons and satellite cells. PTEN expression is markedly reduced in sensory neurons of PTEN iKO mice after 4-HT administration. Scale bar: 50 μm. (B) Western blot analysis of intact WT and PTEN iKO DRG lysates confirms loss of PTEN and activation of PI3K-Akt-mTOR signaling. Molecular mass is indicated in kilodaltons. Spinal cord cross sections show GFP-labeled regenerating axons in WT (C-C’’) and PTEN iKO (D-D’’) mice. GFAP staining denotes the astrocytic boundary of the CNS. Dotted lines indicate the approximate DREZ. Scale bar: 100 μm. (D’’’) A few axons in PTEN iKO mice may penetrate deeper into the DREZ or grow along the pial surface, but they do not grow into the spinal cord gray matter. Scale bar: 50 μm. (E) Axon growth analysis shows no significant difference between WT and PTEN iKO groups at any distance from the GFAP border. n = 5 mice per group, at least 5-6 sections per mouse, unpaired t-test. Values represent mean ± SEM.

We next assessed whether PTEN deletion promotes DR axon regeneration across the DREZ. In control animals, injured DR axons regenerated along the DR but failed to penetrate the DREZ (Figure 5C, E). After PTEN deletion, a few axons appeared to approach or cross the GFAP border (Figure 5D-D’’’), but growth was not observed beyond 100 μm central to the outer astrocytic boundary (Figure 5E). By our classification, this limited growth remained within DREZ territory and did not represent intraspinal regeneration. Axon number index did not differ significantly between control and PTEN iKO groups. Thus, despite activation of PI3K-Akt-mTOR signaling, PTEN deletion alone is insufficient to promote regeneration of DR axons into the spinal cord.

### SOCS3 deletion does not promote axon regeneration across the DREZ

JAK-STAT3 signaling has been implicated in sensory axon regeneration after conditioning lesion 43, and deletion of SOCS3, a negative regulator of JAK-STAT3 signaling, enhances optic nerve regeneration 16. We therefore asked whether SOCS3 deletion in adult sensory neurons improves regeneration of injured DR axons into the spinal cord. We used a brn3aCreERT2:tdTomato:SOCS3f/f transgenic line to conditionally delete SOCS3 from adult DRG sensory neurons. Western blotting showed upregulation of survivin, a downstream pSTAT3 target, consistent with activation of JAK-STAT3 signaling (Figure 6A). As expected, injured DR axons in the control group regenerated along the DR but did not enter the spinal cord (Figure 6B-B’). Contrary to findings from other injury models, SOCS3 deletion did not improve axon regeneration past the DREZ compared with controls (Figure 6C-C’). Axon index analysis revealed no significant difference between control and SOCS3 iKO groups (Figure 6D). These results indicate that SOCS3 deletion alone is not sufficient to overcome the DREZ barrier.

**Figure 6.**
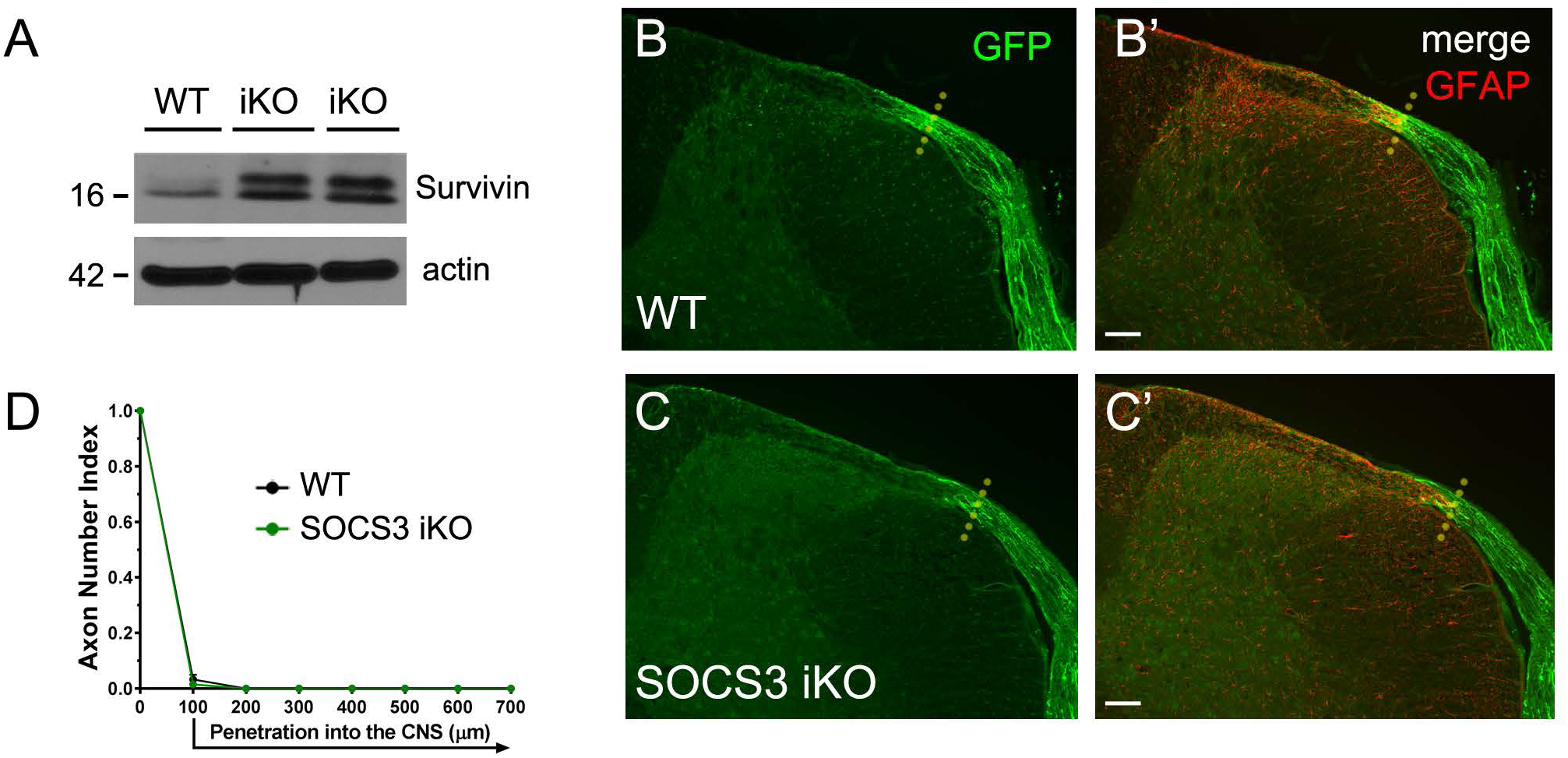
SOCS3 deletion does not promote axon regeneration across the DREZ. (A) Western blotting confirms induction of JAK-STAT3 pathway signaling by upregulation of survivin. GFP-labeled sensory axons are not observed past the DREZ in WT (B-B’) or SOCS3 iKO (C-C’) mice. GFAP staining denotes the astrocytic boundary of the CNS. Dotted lines indicate the approximate DREZ. Scale bar: 100 μm. (D) Axon number index shows no significant difference between groups at any distance. n = 4-5 mice per group, at least 5-6 sections per mouse, unpaired t-test. Values represent mean ± SEM.

### PTEN and SOCS3 co-deletion does not promote axon regeneration across the DREZ

In other CNS injury models, combined deletion of PTEN and SOCS3 produces long-distance regeneration of optic nerve and corticospinal tract axons, likely because PI3K-Akt-mTOR and JAK-STAT3 pathways can act through partially independent mechanisms 15,44. Because PTEN or SOCS3 deletion alone produced little or no regeneration across the DREZ, we tested whether simultaneous deletion of both inhibitors would have a combined effect. In control animals, injured DR axons did not cross the DREZ (Figure 7A-A’). Similarly, in brn3aCreERT2:tdTomato:PTENf/f:SOCS3f/f mice, combined PTEN and SOCS3 deletion failed to promote DR axon regeneration into spinal cord gray matter (Figure 7B-B’). Quantification showed no significant difference in axon growth between the PTEN/SOCS3 iDKO group and SOCS3 iKO, PTEN iKO, or control groups at any distance (Figure 7C). Thus, co-deletion of PTEN and SOCS3 is not sufficient to induce DR axon regeneration across the DREZ.

**Figure 7.**
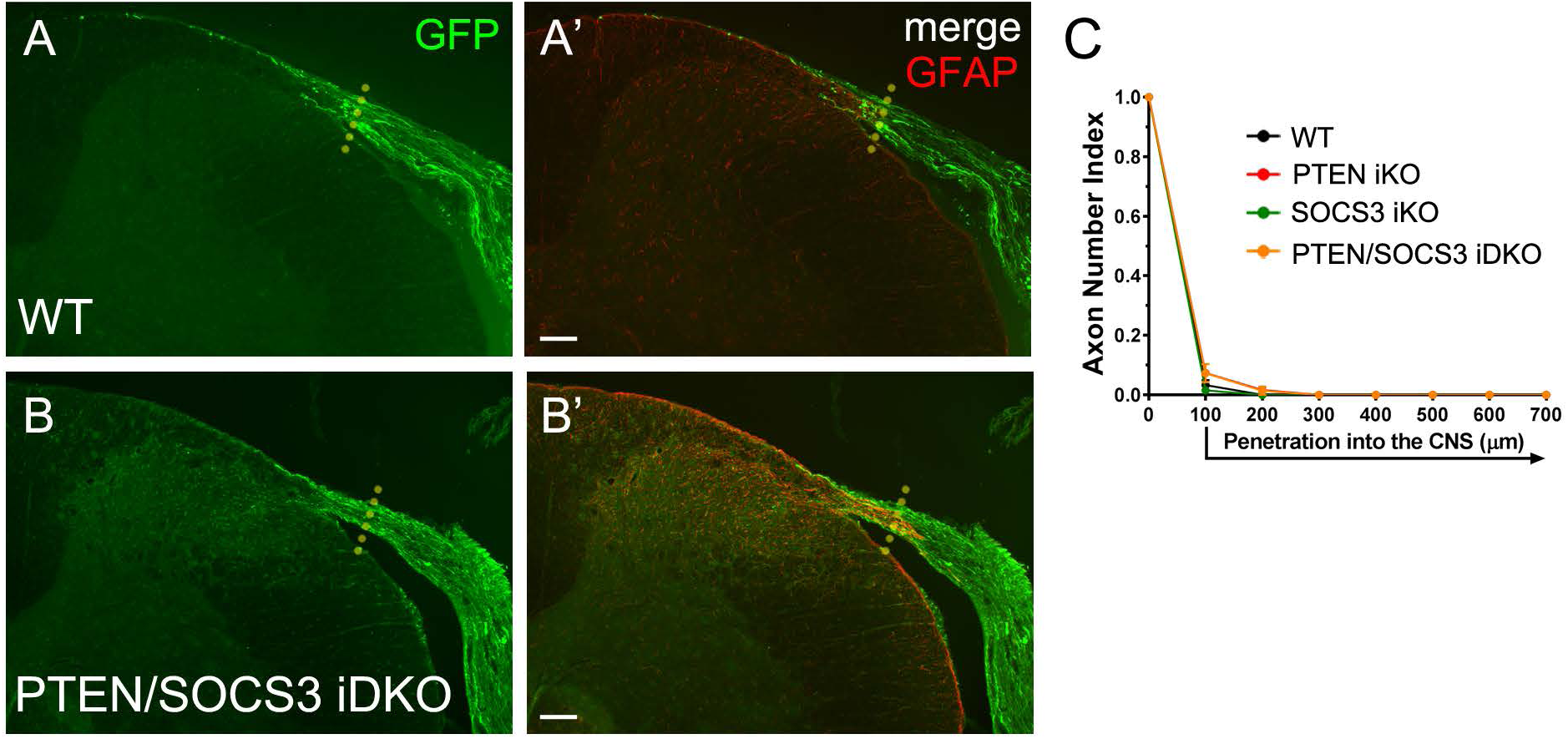
PTEN and SOCS3 co-deletion does not promote intraspinal axon regeneration. Axon regeneration is observed along the DR in both WT (A-A’) and PTEN/SOCS3 iDKO (B-B’) mice, but GFP-labeled axons are not found in the spinal cord in either group. GFAP staining denotes the astrocytic boundary of the CNS. Dotted lines indicate the approximate DREZ. Scale bar: 100 μm. (C) Comparison of axon number index among PTEN/SOCS3 iDKO, SOCS3 iKO, PTEN iKO, and WT groups shows no significant difference at any distance. n = 4-5 mice per group, at least 5-6 sections per mouse, unpaired t-test. Values represent mean ± SEM.

### PTEN deletion markedly enhances kaBRAF-elicited intraspinal regeneration

We next tested whether PTEN deletion could enhance kaBRAF-mediated DR axon regeneration. Using brn3aCreERT2:tdTomato:BRAFV600E:PTENf/f mice, we assessed regeneration 3 weeks after DR injury using the same experimental design (Figure 2A).

Regeneration after PTEN deletion alone (Figure 8A-A’, a) and kaBRAF expression alone (Figure 8B-B’, b) is shown again for comparison. Strikingly, simultaneous kaBRAF expression and PTEN deletion produced robust regeneration of injured DR axons past the DREZ and into the spinal cord (Figure 8C-D). Two major growth patterns were observed. In some animals, axons extended through multiple laminae and deep into the spinal cord, approaching the ventral horn (Figure 8C-C’, c). In others, high-density growth was concentrated in the superficial dorsal horn, with more limited extension into deeper laminae (Figure 8D-D’, d). Despite their increased growth capacity, regenerating axons remained largely confined to gray matter, suggesting that growth was not completely uncontrolled. In wholemount and cross-sectional preparations, axons extended rostrocaudally into cervical levels C2 and C3, showed minor growth into C8, and were not detected in T1 (not shown). Quantification showed that kaBRAF with PTEN deletion significantly increased axon number index at 100, 200, 300, 400, and 500 μm into the spinal cord compared with all other groups (Figure 8E). Axon density was also significantly higher in kaBRAF/PTEN mice than in control, PTEN iKO, or kaBRAF iTg mice (Figure 8F). These data demonstrate that combined BRAF-MEK-ERK activation and PTEN deletion induces robust long-distance intraspinal regeneration within three weeks after injury.

**Figure 8.**
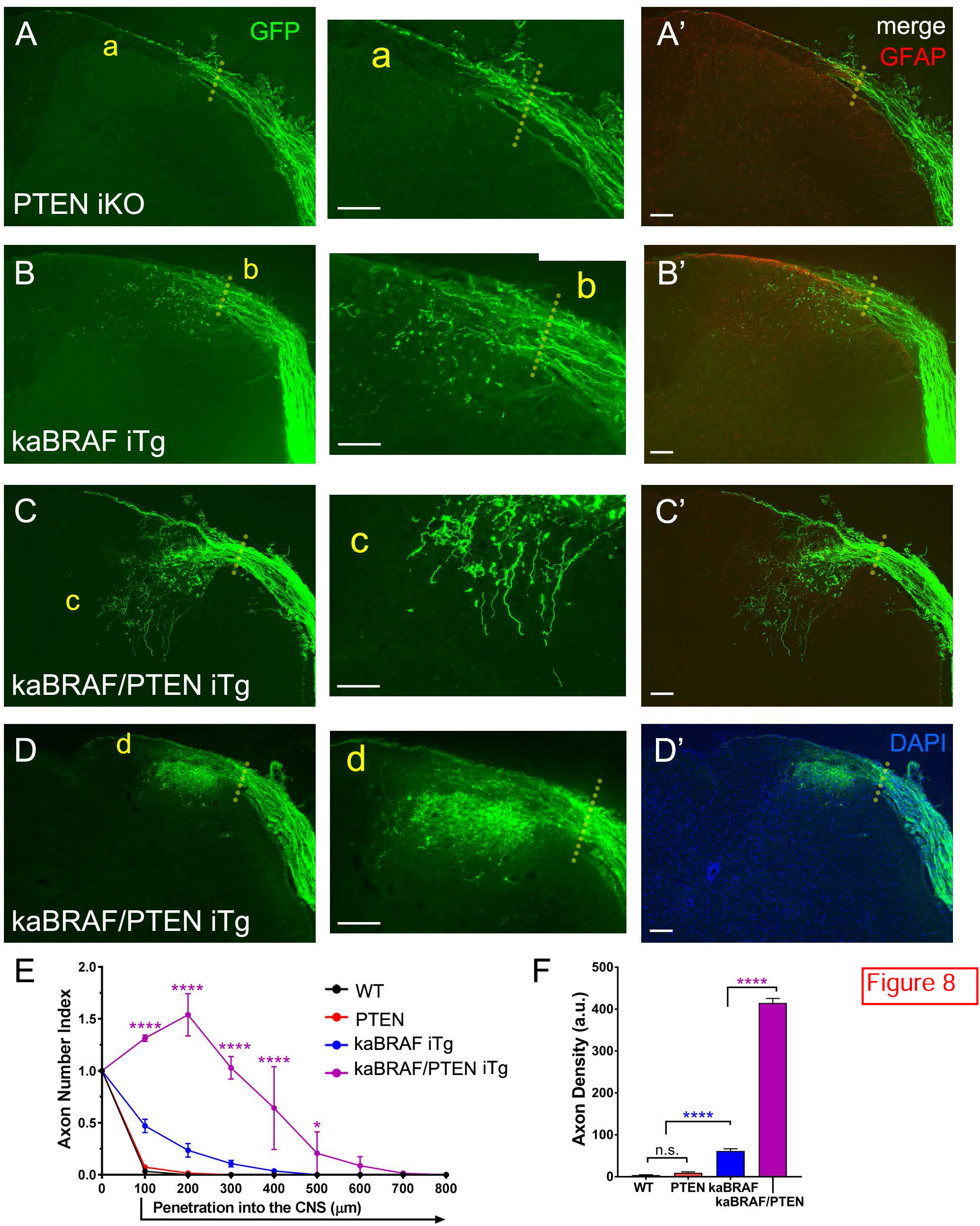
PTEN deletion markedly enhances kaBRAF-elicited intraspinal regeneration. Cross sections show GFP-labeled regenerating axons 3 weeks after DR injury in PTEN iKO (A-A’, a), kaBRAF iTg (B-B’, b), and kaBRAF/PTEN iTg (C-D’, c-d) mice. Simultaneous kaBRAF expression and PTEN deletion induces dramatic axon regeneration into spinal cord gray matter (C-D’). Growth extends deeply toward the ventral horn in one example (c). In another animal of the same genotype, high-density regeneration is concentrated in superficial dorsal horn laminae (d). GFAP staining denotes the astrocytic boundary of the CNS. Dotted lines indicate the approximate DREZ. Scale bars: 100 μm. (E) Quantification of axon growth shows significantly more regeneration in kaBRAF/PTEN iTg mice than in kaBRAF iTg, PTEN iKO, or WT mice at 100, 200, 300, 400, and 500 μm from the GFAP border. Axon number index >1 indicates branching. (F) Axon density is significantly higher in kaBRAF/PTEN iTg mice than in kaBRAF iTg mice. n = 2-5 mice per group, at least 5-6 sections per mouse. **** P < 0.0001, * P < 0.05, n.s. = not significant, one-way or two-way ANOVA with Tukey’s multiple comparisons test.

### SOCS3 deletion does not enhance kaBRAF-elicited intraspinal regeneration

Finally, we examined whether SOCS3 deletion further enhances regeneration in the context of kaBRAF expression or combined kaBRAF expression and PTEN deletion. Using brn3aCreERT2:tdTomato:BRAFV600E:SOCS3f/f mice, we found that regeneration in kaBRAF/SOCS3 mice was comparable to that in kaBRAF mice, indicating that SOCS3 deletion does not enhance kaBRAF-mediated growth across the DREZ at 3 weeks after DR crush (Figure 9A-A’, a). We then tested brn3aCreERT2:tdTomato:BRAFV600E:PTENf/f:SOCS3f/f mice to determine whether simultaneous activation of BRAF-MEK-ERK, PI3K-Akt-mTOR, and JAK-STAT3 signaling would maximize growth competence. Addition of SOCS3 deletion produced only a minor improvement over kaBRAF/PTEN (Figure 9B-C). Axon number index was significantly higher in kaBRAF/PTEN/SOCS3 mice than in kaBRAF/PTEN mice at 600 μm from the DREZ, but no significant differences were detected at most other distances (Figure 9D). Axon density analysis showed only a small increase in overall growth in kaBRAF/PTEN/SOCS3 mice compared with kaBRAF/PTEN mice (Figure 9E). Thus, concurrent activation of BRAF-MEK-ERK and PI3K-Akt-mTOR signaling is the key driver of sustained long-distance DR axon regeneration, whereas JAK-STAT3 activation through SOCS3 deletion provides little additional benefit.

**Figure 9.**
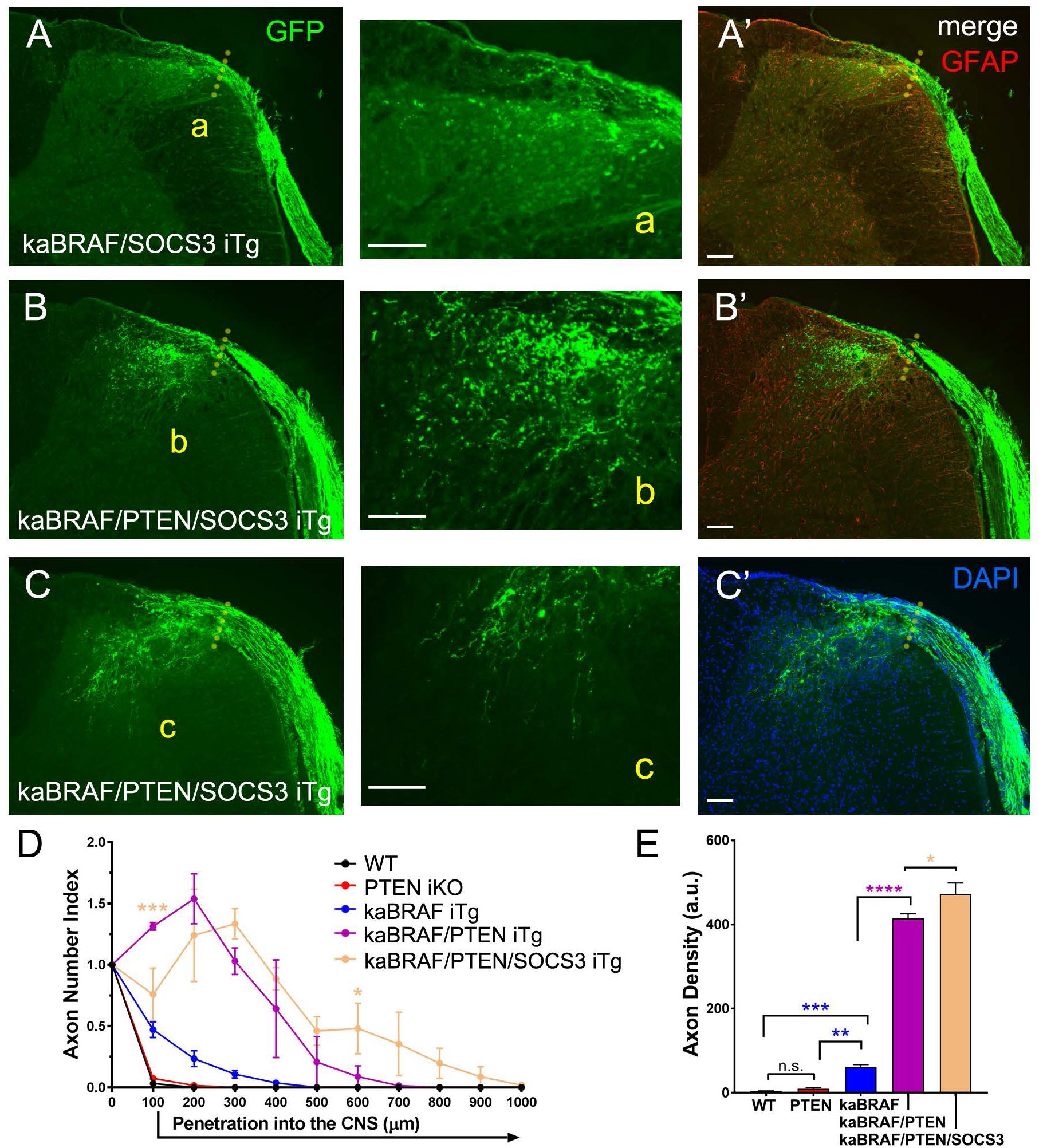
SOCS3 deletion does not enhance kaBRAF-elicited intraspinal regeneration. (A-A’, a) kaBRAF expression with SOCS3 deletion promotes regeneration across the DREZ at 3 weeks after injury, but the extent of growth is comparable to kaBRAF expression alone (quantitative data not shown). (B-C’) Three weeks after DR injury, robust GFP-labeled sensory axon regeneration is observed in kaBRAF/PTEN/SOCS3 iTg mice. Two examples are shown. (b-c) Axons penetrate the DREZ and grow deeply into the spinal cord, but the overall response is comparable to kaBRAF/PTEN iTg mice. GFAP staining denotes the astrocytic boundary of the CNS. Dotted lines indicate the approximate DREZ. Scale bars: 100 μm. (D) Axon number index shows a significant difference between kaBRAF/PTEN/SOCS3 iTg and kaBRAF/PTEN iTg groups at 100 μm and 600 μm. Reduced growth at 100 μm in the kaBRAF/PTEN/SOCS3 iTg group is likely attributable to inflammatory disruption near the DREZ in some animals. (E) Axon density analysis shows a slight but significant difference between kaBRAF/PTEN/SOCS3 iTg and kaBRAF/PTEN iTg groups. n = 2-5 mice per group, at least 5-6 sections per mouse. **** P < 0.0001, *** P < 0.001, ** P < 0.01, * P < 0.05, n.s. = not significant, one-way or two-way ANOVA with Tukey’s multiple comparisons test.

## DISCUSSION

This study identifies combined activation of BRAF-MEK-ERK and PI3K-Akt-mTOR signaling as a powerful strategy for promoting intraspinal regeneration of injured DR axons. Constitutively active B-RAF or PTEN deletion alone produced only limited axon entry into, or near, the DREZ. In contrast, simultaneous kaBRAF expression and PTEN deletion induced extensive growth deep into the spinal cord, indicating a strong cooperative effect between these two intrinsic growth pathways. SOCS3 deletion, either alone or in combination with PTEN deletion or kaBRAF expression, did not substantially enhance regeneration. Likewise, removal of the myelin-associated inhibitors Nogo, MAG, and OMgp produced only modest additional benefit when combined with kaBRAF. Together, these findings argue that the intrinsic growth state of adult sensory neurons is a dominant determinant of regeneration across the DREZ and that dual engagement of RAF-MEK-ERK and PI3K-Akt-mTOR signaling is particularly effective in large-diameter DR axons.

A key motivation for this work was to re-evaluate kaBRAF-mediated regeneration under conditions that minimize confounding inflammatory lesions near the DREZ. In our previous study, kaBRAF expression was associated with significant growth into superficial dorsal horn laminae, but the presence of inflammatory disruption at the DREZ complicated interpretation.

Lesion-associated cells and inflammatory microenvironments can alter sensory axon behavior and may provide growth-permissive substrates under some conditions 76. Here, by excluding spinal cords with overt disruption of the GFAP+ border, we confirmed that kaBRAF itself promotes reproducible axon entry across the DREZ, although the effect is limited. This result is consistent with the established role of B-RAF in sensory neuron survival and developmental sensory axon growth 21,22,45, and extends previous evidence that constitutive B-RAF activation increases neurite outgrowth in DRG neurons and promotes regeneration in the mature CNS 23.

Continued kaBRAF expression sustained limited anatomical growth over two months, but this was not accompanied by significant sensory recovery. This dissociation is important because it reinforces the idea that anatomical growth alone is not sufficient for functional repair; regenerating axons must also reach appropriate target regions, form synapses, and integrate into functional circuits 80.

PTEN deletion alone also failed to promote meaningful intraspinal regeneration despite clear activation of PI3K-Akt-mTOR signaling, as indicated by pS6 upregulation. This result contrasts with robust effects of PTEN deletion in retinal ganglion cells and corticospinal neurons, where PTEN loss enhances optic nerve and corticospinal tract regeneration 17,41,46–49. Our findings do not imply that PI3K-Akt-mTOR signaling is irrelevant in adult sensory neurons.

Rather, they suggest that PTEN deletion alone is insufficient to overcome the combined intrinsic and extrinsic constraints imposed at the DREZ. In cultured NF+ DRG neurons, pharmacological PTEN inhibition promotes neurite outgrowth 42,77, supporting the idea that PI3K-Akt signaling can enhance sensory axon growth under permissive conditions. In vivo, however, the DREZ appears to require stronger or broader growth activation than PTEN deletion alone can provide. The striking effect of kaBRAF/PTEN therefore suggests that PI3K-Akt-mTOR signaling becomes most effective when paired with a second pathway that drives a complementary regenerative program.

In contrast to PTEN, SOCS3 deletion had little effect in this model. This was unexpected because JAK-STAT3 signaling is required for conditioning lesion-induced sensory axon regeneration in the dorsal column 43,50–52, and SOCS3 deletion promotes optic nerve regeneration 15,16. However, our findings are consistent with in vitro studies showing that SOCS3 manipulation has limited effects on unconditioned DRG neurons and becomes more influential after a prior peripheral nerve injury 53. One limitation is that we did not administer exogenous cytokines such as CNTF, which can potentiate SOCS3-dependent regeneration and pSTAT3 activation in the optic nerve 15,16,53. Thus, we cannot exclude the possibility that SOCS3 deletion might enhance DR regeneration under conditions that provide stronger cytokine stimulation or conditioning lesion-like signals. Nevertheless, under the conditions tested here, SOCS3 deletion was neither sufficient by itself nor substantially additive to kaBRAF/PTEN-mediated growth.

Our analysis of myelin-associated inhibitors further supports the view that reducing extrinsic inhibition alone is not sufficient to promote robust DR regeneration. Deletion of Nogo, MAG, and OMgp did not enhance regeneration across the DREZ compared with controls, and its combination with kaBRAF produced only a modest increase. This finding is consistent with previous studies showing limited regeneration in mice lacking major myelin-associated inhibitors 39,54 and with work showing that Nogo deletion does not markedly augment PTEN-driven corticospinal regeneration 55. These results do not exclude a role for extrinsic inhibitors at the DREZ. Instead, they suggest that, in adult sensory neurons, overcoming myelin-associated inhibition requires a sufficiently strong intrinsic growth state. In this context, kaBRAF appears to provide a partial growth stimulus, whereas combined kaBRAF/PTEN produces a much stronger response.

The most important finding of this study is the synergistic effect of kaBRAF expression and PTEN deletion. kaBRAF/PTEN increased both axon number and axon density within the spinal cord, suggesting that the two pathways may act on distinct but complementary aspects of regeneration. One possibility is that BRAF-MEK-ERK signaling promotes transcriptional and biosynthetic programs required for axon elongation, whereas PI3K-Akt-mTOR signaling enhances growth cone dynamics, cytoskeletal remodeling, and local protein synthesis. Dual pathway activation may therefore generate a broader and more sustained regenerative state than either pathway alone 15,57,58. Mechanistically, PTEN deletion can promote GSK3β inactivation 59, Rac/Cdc42-dependent filopodia and lamellipodia formation 60, increased growth cone responsiveness to chemoattractive cues 61, and improved injury signaling through ERK-related pathways 52,58,62,63. In parallel, kaBRAF activation in the cell body may support regeneration-associated gene expression through sustained ERK signaling and altered feedback regulation of RAF-MEK-ERK pathway output 18,52,57,64,65. The oncogenic origin of BRAFV600E 78 and the tumor-suppressor role of PTEN also underscore the need for caution in translating these findings. Future work should test whether temporally controlled, non-genetic, or locally targeted approaches can reproduce the regenerative benefit while reducing potential safety concerns.

The present results place kaBRAF/PTEN-mediated intrinsic activation among the most effective strategies for promoting regeneration of large-diameter DR axons across the DREZ. Conditioning lesion paradigms 27,66, neurotrophic factor delivery or expression 12-14,31,67-70, cAMP elevation 73,74, and transcription factor-based approaches 71,72 have all provided important evidence that adult DRG neurons can be pushed toward a regenerative state. However, neurotrophic factor-based strategies often act selectively on receptor-defined sensory subtypes and can promote ectopic growth or inappropriate laminar targeting when expressed locally in the spinal cord 14,75. Direct activation of downstream intrinsic signaling pathways may offer a complementary strategy that avoids overwhelming the intraspinal environment with exogenous trophic cues. At the same time, our behavioral data and the broader field emphasize that regeneration must be anatomically precise and functionally integrated, not simply extensive 80. Future studies should determine whether kaBRAF/PTEN-regenerated axons form synapses with appropriate intraspinal targets, whether small-diameter sensory axons can be similarly recruited, and whether pathway activation after injury can produce functional recovery. Overall, our findings identify combined BRAF-MEK-ERK and PI3K-Akt-mTOR activation as a potent strategy for overcoming the DREZ barrier and promoting long-distance regeneration of injured sensory axons.

## Acknowledgements

We thank members of the Son laboratory for critical reading of the manuscript. We thank Dr. Huaqing Zhao for statistical help and Drs. Matthew Grove and Gareth Thomas for assistance with Western blotting. This work was supported by Shriners Hospitals for Children and NIH NINDS (NS079631 and NS095070 to Y.-J.S.).

## REFERENCES

1. Kaiser, R., Waldauf, P., Ullas, G., and Krajcová, A. (2020). Epidemiology, etiology, and types of severe adult brachial plexus injuries requiring surgical repair: systematic review and meta-analysis. Neurosurg Rev 43, 443–452. 10.1007/s10143-018-1009-2.

2. Davis, G., and Curtin, C.M. (2016). Management of Pain in Complex Nerve Injuries. Hand Clinics 32, 257-+. 10.1016/j.hcl.2015.12.011.

3. Smania, N., Berto, G., La Marchina, E., Melotti, C., Midiri, A., Roncari, L., Zenorini, A., Ianes, P., Picelli, A., Waldner, A., et al. (2012). Rehabilitation of brachial plexus injuries in adults and children. European Journal of Physical and Rehabilitation Medicine 48, 483–506.

4. Caroni, P., Savio, T., and Schwab, M.E. (1988). Central nervous system regeneration - oligodendrocytes and myelin as non-permissive substrates for neurite growth. Progress in Brain Research 78, 363–370. 10.1016/s0079-6123(08)60305-2.

5. Schnell, L., and Schwab, M.E. (1990). Axonal regeneration in the rat spinal cord produced by an antibody against myelin-associated neurite growth-inhibitors. Nature 343, 269–272. 10.1038/343269a0.

6. McKeon, R.J., Hoke, A., and Silver, J. (1995). Injury-induced proteoglycans inhibit the potential for laminin-mediated axon growth on astrocytic scars. Experimental Neurology 136, 32–43. 10.1006/exnr.1995.1081.

7. McKeon, R.J., Schreiber, R.C., Rudge, J.S., and Silver, J. (1991). Reduction of neurite outgrowth in a model of glial scarring following CNS injury is correlated with the expression of inhibitory molecules on reactive astrocytes. Journal of Neuroscience 11, 3398–3411.

8. Cafferty, W.B.J., Bradbury, E.J., Lidierth, M., Jones, M., Duffy, P.J., Pezet, S., and McMahon, S.B. (2008). Chondroitinase ABC-Mediated Plasticity of Spinal Sensory Function. Journal of Neuroscience 28, 11998–12009. 10.1523/jneurosci.3877-08.2008.

9. Golding, J.P., Bird, C., McMahon, S., and Cohen, J. (1999). Behaviour of DRG sensory neurites at the intact and injured adult rat dorsal root entry zone: postnatal neurites become paralysed, whilst injury improves the growth of embryonic neurites. Glia 26, 309–323. 10.1002/(SICI)1098-1136(199906)26:4<309::AID-GLIA5>3.0.CO;2-0 [pii].

10. Golding, J.P., Shewan, D., Berry, M., and Cohen, J. (1996). An in vitro model of the rat dorsal root entry zone reveals developmental changes in the extent of sensory axon growth into the spinal cord. Molecular and Cellular Neuroscience 7, 191–203. 10.1006/mcne.1996.0015.

11. Lutz, A.B., and Barres, B.A. (2014). Contrasting the Glial Response to Axon Injury in the Central and Peripheral Nervous Systems. Developmental cell 28, 7–17. 10.1016/j.devcel.2013.12.002.

12. Ramer, M.S., Duraisingam, I., Priestley, J.V., and McMahon, S.B. (2001). Two-tiered inhibition of axon regeneration at the dorsal root entry zone. Journal of Neuroscience 21, 2651–2660.

13. Ramer, M.S., Priestley, J.V., and McMahon, S.B. (2000). Functional regeneration of sensory axons into the adult spinal cord. Nature 403, 312–316. 10.1038/35002084.

14. Kelamangalath, L., Tang, X., Bezik, K., Sterling, N., Son, Y.J., and Smith, G.M. (2015). Neurotrophin selectivity in organizing topographic regeneration of nociceptive afferents. Experimental Neurology 6, 262–278.

15. Sun, F., Park, K.K., Belin, S., Wang, D., Lu, T., Chen, G., Zhang, K., Yeung, C., Feng, G., Yankner, B.A., and He, Z. (2011). Sustained axon regeneration induced by co-deletion of PTEN and SOCS3. Nature 480, 372–U125. 10.1038/nature10594.

16. Smith, P.D., Sun, F., Park, K.K., Cai, B., Wang, C., Kuwako, K., Martinez-Carrasco, I., Connolly, L., and He, Z. (2009). SOCS3 Deletion Promotes Optic Nerve Regeneration In Vivo. Neuron 64, 617–623. 10.1016/j.neuron.2009.11.021.

17. Park, K.K., Liu, K., Hu, Y., Smith, P.D., Wang, C., Cai, B., Xu, B.G., Connolly, L., Kramvis, I., Sahin, M., and He, Z.G. (2008). Promoting Axon Regeneration in the Adult CNS by Modulation of the PTEN/mTOR Pathway. Science 322, 963–966. 10.1126/science.1161566.

18. Mihaly, A., Priestley, J.V., and Molnar, E. (1996). Expression of raf serine/threonine protein kinases in the cell bodies of primary sensory neurons of the adult rat. Cell and Tissue Research 285, 261–271. 10.1007/s004410050643.

19. Roberts, P.J., and Der, C.J. (2007). Targeting the Raf-MEK-ERK mitogen-activated protein kinase cascade for the treatment of cancer. Oncogene 26, 3291–3310. 10.1038/sj.onc.1210422.

20. Matallanas, D., Birtwistle, M., Romano, D., Zebisch, A., Rauch, J., von Kriegsheim, A., and Kolch, W. (2011). Raf Family Kinases: Old Dogs have Learned New Tricks. Genes & Cancer 2, 232–260.

21. Markus, A., Zhong, J., and Snider, W.D. (2002). Raf and akt mediate distinct aspects of sensory axon growth. Neuron 35, 65–76.

22. Zhong, J., Li, X., McNamee, C., Chen, A.P., Baccarini, M., and Snider, W.D. (2007). Raf kinase signaling functions in sensory neuron differentiation and axon growth in vivo. Nat. Neurosci. 10, 598–607. 10.1038/1898.

23. O’Donovan, K.J., Ma, K., Guo, H., Wang, C., Sun, F., Han, S.B., Kim, H., Wong, J.K., Charron, J., Zou, H., et al. (2014). B-RAF kinase drives developmental axon growth and promotes axon regeneration in the injured mature CNS. Journal of Experimental Medicine 211, 801–814. 10.1084/jem.20131780.

24. Eng, S.R., Gratwick, K., Rhee, J.M., Fedtsova, N., Gan, L., and Turner, E.E. (2001). Defects in sensory axon growth precede neuronal death in Brn3a-deficient mice. Journal of Neuroscience 21, 541–549.

25. Mercer, K., Giblett, S., Green, S., Lloyd, D., Dias, S.D., Plumb, M., Marais, R., and Pritchard, C. (2005). Expression of endogenous oncogenic B-V600E-raf induces proliferation and developmental defects in mice and transformation of primary fibroblasts. Cancer Research 65, 11493–11500. 10.1158/0008-5472.can-05-2211.

26. Liu, Y.P., Kelamangalath, L., Kim, H., Han, S.B., Tang, X.Q., Zhai, J.B., Hong, J.W., Lin, S., Son, Y.J., and Smith, G.M. (2016). NT-3 promotes proprioceptive axon regeneration when combined with activation of the mTor intrinsic growth pathway but not with reduction of myelin extrinsic inhibitors. Experimental neurology 283, 73–84. 10.1016/j.expneurol.2016.05.021.

27. Di Maio, A., Skuba, A., Himes, B.T., Bhagat, S.L., Hyun, J.K., Tessler, A., Bishop, D., and Son, Y.J. (2011). In Vivo Imaging of Dorsal Root Regeneration: Rapid Immobilization and Presynaptic Differentiation at the CNS/PNS Border. The Journal of neuroscience : the official journal of the Society for Neuroscience 31, 4569–4582. 31/12/4569 [pii] 10.1523/JNEUROSCI.4638-10.2011.

28. Steward, O., Zheng, B., Tessier-Lavigne, M., Hofstadter, M., Sharp, K., and Yee, K.M. (2008). Regenerative growth of corticospinal tract axons via the ventral column after spinal cord injury in mice. Journal of Neuroscience 28, 6836–6847. 10.1523/jneurosci.5372-07.2008.

29. Ma, M.H., Basso, D.M., Walters, P., Stokes, B.T., and Jakeman, L.B. (2001). Behavioral and histological outcomes following graded spinal cord contusion injury in the C57Bl/6 mouse. Experimental neurology 169, 239–254. 10.1006/exnr.2001.7679.

30. Metz, G.A., and Whishaw, I.Q. (2002). Cortical and subcortical lesions impair skilled walking in the ladder rung walking test: a new task to evaluate fore- and hindlimb stepping, placing, and co-ordination. Journal of Neuroscience Methods 115, 169–179, Pii s0165-0270(02)00012-2. 10.1016/s0165-0270(02)00012-2.

31. Wang, R., King, T., Ossipov, M.H., Rossomando, A.J., Vanderah, T.W., Harvey, P., Cariani, P., Frank, E., Sah, D.W.Y., and Porreca, F. (2008). Persistent restoration of sensory function by immediate or delayed systemic artemin after dorsal root injury. Nature Neuroscience 11, 488–496. 10.1038/nn2069.

32. Chaplan, S.R., Bach, F.W., Pogrel, J.W., Chung, J.M., and Yaksh, T.L. (1994). Quantitative assessment of tactile allodynia in the rat paw. Journal of Neuroscience Methods 53, 55–63. 10.1016/0165-0270(94)90144-9.

33. Dixon, W.J. (1980). Efficient analysis of experimental observations. Annual Review of Pharmacology and Toxicology 20, 441–462. 10.1146/annurev.pa.20.040180.002301.

34. Mogil, J.S., Wilson, S.G., Bon, K., Lee, S.E., Chung, K., Raber, P., Pieper, J.O., Hain, H.S., Belknap, J.K., Hubert, L., et al. (1999). Heritability of nociception I: Responses of 11 inbred mouse strains on 12 measures of nociception. Pain 80, 67–82. 10.1016/s0304-3959(98)00197-3.

35. Grider, M.H., Chen, Q., and Shine, H.D. (2006). Semi-automated quantification of axonal densities in labeled CNS tissue. Journal of Neuroscience Methods 155, 172–179. 10.1016/j.jneumeth.2005.12.021.

36. Liu, Y.P., Keefe, K., Tang, X.Q., Lin, S., and Smith, G.M. (2014). Use of Self-Complementary Adeno-Associated Virus Serotype 2 as a Tracer for Labeling Axons: Implications for Axon Regeneration. PloS one 9, e87447. 10.1371/journal.pone.0087447.

37. Aschauer, D.F., Kreuz, S., and Rumpel, S. (2013). Analysis of Transduction Efficiency, Tropism and Axonal Transport of AAV Serotypes 1, 2, 5, 6, 8 and 9 in the Mouse Brain. PloS one 8, UNSP e76310. 10.1371/journal.pone.0076310.

38. Xiao, X., Li, J., McCown, T.J., and Samulski, R.J. (1997). Gene transfer by adeno-associated virus vectors into the central nervous system. Experimental neurology 144, 113–124. 10.1006/exnr.1996.6396.

39. Lee, J.K., Geoffroy, C.G., Chan, A.F., Tolentino, K.E., Crawford, M.J., Leal, M.A., Kang, B., and Zheng, B.H. (2010). Assessing Spinal Axon Regeneration and Sprouting in Nogo-, MAG-, and OMgp-Deficient Mice. Neuron 66, 663–670. 10.1016/j.neuron.2010.05.002.

40. Park, K.K., Liu, K., Hu, Y., Kanter, J.L., and He, Z. (2010). PTEN/mTOR and axon regeneration. Experimental Neurology 223, 45–50. 10.1016/j.expneurol.2009.12.032.

41. Liu, K., Lu, Y., Lee, J.K., Samara, R., Willenberg, R., Sears-Kraxberger, I., Tedeschi, A., Park, K.K., Jin, D., Cai, B., et al. (2010). PTEN deletion enhances the regenerative ability of adult corticospinal neurons. Nat. Neurosci. 13, 1075–U1064. 10.1038/nn.2603.

42. Christie, K.J., Webber, C.A., Martinez, J.A., Singh, B., and Zochodne, D.W. (2010). PTEN Inhibition to Facilitate Intrinsic Regenerative Outgrowth of Adult Peripheral Axons. Journal of Neuroscience 30, 9306–9315. 10.1523/jneurosci.6271-09.2010.

43. Qiu, J., Cafferty, W.B.J., McMahon, S.B., and Thompson, S.W.N. (2005). Conditioning injury-induced spinal axon regeneration requires signal transducer and activator of transcription 3 activation. Journal of Neuroscience 25, 1645–1653. 10.1523/jneurosci.3269-04.2005.

44. Jin, D., Liu, Y.Y., Sun, F., Wang, X.H., Liu, X.F., and He, Z.G. (2015). Restoration of skilled locomotion by sprouting corticospinal axons induced by co-deletion of PTEN and SOCS3. Nature Communications 6, 8074. 10.1038/ncomms9074.

45. Wiese, S., Pei, G., Karch, C., Troppmair, J., Holtmann, B., Rapp, U.R., and Sendtner, M. (2001). Specific function of B-RAF in mediating survival of embryonic motoneurons and sensory neurons. Nature Neuroscience 4, 137–142. 10.1038/83960.

46. Duan, X., Qiao, M., Bei, F.F., Kim, I.J., He, Z.G., and Sanes, J.R. (2015). Subtype-Specific Regeneration of Retinal Ganglion Cells following Axotomy: Effects of Osteopontin and mTOR Signaling. Neuron 85, 1244–1256. 10.1016/j.neuron.2015.02.017.

47. Danilov, C.A., and Steward, O. (2015). Conditional genetic deletion of PTEN after a spinal cord injury enhances regenerative growth of CST axons and motor function recovery in mice. Experimental neurology 266, 147–160. 10.1016/j.expneurol.2015.02.012.

48. Du, K.M., Zheng, S.S., Zhang, Q., Li, S.S., Gao, X., Wang, J., Jiang, L.W., and Liu, K. (2015). Pten Deletion Promotes Regrowth of Corticospinal Tract Axons 1 Year after Spinal Cord Injury. Journal of Neuroscience 35, 9754–9763. 10.1523/jneurosci.3637-14.2015.

49. Zukor, K., Belin, S., Wang, C., Keelan, N., Wang, X.H., and He, Z.G. (2013). Short Hairpin RNA against PTEN Enhances Regenerative Growth of Corticospinal Tract Axons after Spinal Cord Injury. Journal of Neuroscience 33, 15350–15361. 10.1523/jneurosci.2510-13.2013.

50. Cafferty, W.B.J., Gardiner, N.J., Das, P., Qiu, J., McMahon, S.B., and Thompson, S.W.N. (2004). Conditioning injury-induced spinal axon regeneration fails in interleukin-6 knock-out mice. Journal of Neuroscience 24, 4432–4443. 10.1523/jneurosci.2245-02.2004.

51. Cafferty, W.B.J., Gardiner, N.J., Gavazzi, I., Powell, J., McMahon, S.B., Heath, J.K., Munson, J., Cohen, J., and Thompson, S.W.N. (2001). Leukemia inhibitory factor determines the growth status of injured adult sensory neurons. Journal of Neuroscience 21, 7161–7170.

52. Zhou, F.Q., and Snider, W.D. (2006). Intracellular control of developmental and regenerative axon growth. Philosophical Transactions of the Royal Society B-Biological Sciences 361, 1575–1592. 10.1098/rstb.2006.1882.

53. Miao, T., Wu, D., Zhang, Y., Bo, X., Subang, M.C., Wang, P., and Richardson, P.M. (2006). Suppressor of cytokine signaling-3 suppresses the ability of activated signal transducer and activator of transcription-3 to stimulate neurite growth in rat primary sensory neurons. Journal of Neuroscience 26, 9512–9519. 10.1523/jneurosci.2160-06.2006.

54. Lee, J.K., Chan, A.F., Luu, S.M., Zhu, Y.H., Ho, C., Tessier-Lavigne, M., and Zheng, B.H. (2009). Reassessment of Corticospinal Tract Regeneration in Nogo-Deficient Mice. Journal of Neuroscience 29, 8649–8654. 10.1523/jneurosci.1864-09.2009.

55. Geoffroy, C.G., Lorenzana, A.O., Kwan, J.P., Lin, K., Ghassemi, O., Ma, A., Xu, N., Creger, D., Liu, K., He, Z.G., and Zheng, B.H. (2015). Effects of PTEN and Nogo Codeletion on Corticospinal Axon Sprouting and Regeneration in Mice. Journal of Neuroscience 35, 6413–6428. 10.1523/jneurosci.4013-14.2015.

56. Luo, X.T., Ribeiro, M., Bray, E.R., Lee, D.H., Yungher, B.J., Mehta, S.T., Thakor, K.A., Diaz, F., Lee, J.K., Moraes, C.T., et al. (2016). Enhanced Transcriptional Activity and Mitochondrial Localization of STAT3 Co-induce Axon Regrowth in the Adult Central Nervous System. Cell reports 15, 398–410. 10.1016/j.celrep.2016.03.029.

57. Andreadi, C., Noble, C., Patel, B., Jin, H., Hernandez, M.M.A., Balmanno, K., Cook, S.J., and Pritchard, C. (2012). Regulation of MEK/ERK pathway output by subcellular localization of B-RAF. Biochemical Society Transactions 40, 67–72. 10.1042/bst20110621.

58. Gu, J.G., Tamura, M., and Yamada, K.M. (1998). Tumor suppressor PTEN inhibits integrin- and growth factor-mediated mitogen-activated protein (MAP) kinase signaling pathways. Journal of Cell Biology 143, 1375–1383. 10.1083/jcb.143.5.1375.

59. Zhou, F.Q., Zhou, J., Dedhar, S., Wu, Y.H., and Snider, W.D. (2004). NGF-induced axon growth is mediated by localized inactivation of GSK-30 and functions of the microtubule plus end binding protein APC. Neuron 42, 897–912. 10.1016/j.neuron.2004.05.011.

60. Dent, E.W., Gupton, S.L., and Gertler, F.B. (2011). The growth cone cytoskeleton in axon outgrowth and guidance. Cold Spring Harbor perspectives in biology 3, a001800.

61. Henle, S.J., Carlstrom, L.P., Cheever, T.R., and Henley, J.R. (2013). Differential Role of PTEN Phosphatase in Chemotactic Growth Cone Guidance. Journal of Biological Chemistry 288, 20837–20842. 10.1074/jbc.C113.487066.

62. Atwal, J.K., Singh, K.K., Tessier-Lavigne, M., Miller, F.D., and Kaplan, D.R. (2003). Semaphorin 3F antagonizes neurotrophin-induced phosphatidylinositol 3-kinase and mitogen-activated protein kinase kinase signaling: A mechanism for growth cone collapse. Journal of Neuroscience 23, 7602–7609.

63. Perlson, E., Hanz, S., Ben-Yaakov, K., Segal-Ruder, Y., Seger, R., and Fainzilber, M. (2005). Vimentin-dependent spatial translocation of an activated MAP kinase in injured nerve. Neuron 45, 715–726. 10.1016/j.neuron.2005.01.023.

64. Ginty, D.D., and Segal, R.A. (2002). Retrograde neurotrophin signaling: Trk-ing along the axon. Current opinion in neurobiology 12, 268–274. 10.1016/s0959-4388(02)00326-4.

65. Ritt, D.A., Monson, D.M., Specht, S.I., and Morrison, D.K. (2010). Impact of Feedback Phosphorylation and Raf Heterodimerization on Normal and Mutant B-RAF Signaling. Molecular and Cellular Biology 30, 806–819. 10.1128/mcb.00569-09.

66. Chong, M.S., Woolf, C.J., Haque, N.S.K., and Anderson, P.N. (1999). Axonal regeneration from injured dorsal roots into the spinal cord of adult rats. J. Comp. Neurol. 410, 42–54.

67. Oudega, M., and Hagg, T. (1996). Nerve growth factor promotes regeneration of sensory axons into adult rat spinal cord. Experimental neurology 140, 218–229. 10.1006/exnr.1996.0131.

68. Ramer, M.S., Bishop, T., Dockery, P., Mobarak, M.S., O’Leary, D., Fraher, J.P., Priestley, J.V., and McMahon, S.B. (2002). Neurotrophin-3-mediated regeneration and recovery of proprioception following dorsal rhizotomy. Molecular and Cellular Neuroscience 19, 239–249. 10.1006/mcne.2001.1067.

69. Romero, M.I., Rangappa, N., Garry, M.G., and Smith, G.M. (2001). Functional regeneration of chronically injured sensory afferents into adult spinal cord after neurotrophin gene therapy. Journal of Neuroscience 21, 8408–8416.

70. Zhang, Y., Dijkhuizen, P.A., Anderson, P.N., Lieberman, A.R., and Verhaagen, J. (1998). NT-3 delivered by an adenoviral vector induces injured dorsal root axons to regenerate into the spinal cord of adult rats. Journal of Neuroscience Research 54, 554–562.

71. Cai, D.M., Shen, Y.J., De Bellard, M., Tang, S., and Filbin, M.T. (1999). Prior exposure to neurotrophins blocks inhibition of axonal regeneration by MAG and myelin via a cAMP-dependent mechanism. Neuron 22, 89–101. 10.1016/s0896-6273(00)80681-9.

72. Fagoe, N.D., Attwell, C.L., Kouwenhoven, D., Verhaagen, J., and Mason, M.R.J. (2015). Overexpression of ATF3 or the combination of ATF3, c-Jun, STAT3 and Smad1 promotes regeneration of the central axon branch of sensory neurons but without synergistic effects. Human molecular genetics 24, 6788–6800. 10.1093/hmg/ddv383.

73. Neumann, S., Bradke, F., Tessier-Lavigne, M., and Basbaum, A.I. (2002). Regeneration of sensory axons within the injured spinal cord induced by intraganglionic cAMP elevation. Neuron 34, 885–893. S089662730200702X [pii].

74. Qiu, J., Cai, C.M., Dai, H.N., McAtee, M., Hoffman, P.N., Bregman, B.S., and Filbin, M.T. (2002). Spinal axon regeneration induced by elevation of cyclic AMP. Neuron 34, 895–903. 10.1016/s0896-6273(02)00730-4.

75. Thoenen, H., and Sendtner, M. (2002). Neurotrophins: from enthusiastic expectations through sobering experiences to rational therapeutic approaches. Nature Neuroscience 5, 1046–1050. 10.1038/nn938.

76. Busch, S.A., Horn, K.P., Cuascut, F.X., Hawthorne, A.L., Bai, L.H., Miller, R.H., and Silver, J. (2010). Adult NG2+ cells are permissive to neurite outgrowth and stabilize sensory axons during macrophage-induced axonal dieback after spinal cord injury. Journal of Neuroscience 30, 255–265. 10.1523/jneurosci.3705-09.2010.

77. Rosivatz, E., Matthews, J.G., McDonald, N.Q., Mulet, X., Ho, K.K., Lossi, N., Schmid, A.C., Mirabelli, M., Pomeranz, K.M., Erneux, C., et al. (2006). A small molecule inhibitor for phosphatase and tensin homologue deleted on chromosome 10 (PTEN). ACS chemical biology 1, 780–790. 10.1021/cb600352f.

78. Davies, H., Bignell, G.R., Cox, C., Stephens, P., Edkins, S., Clegg, S., Teague, J., Woffendin, H., Garnett, M.J., Bottomley, W., et al. (2002). Mutations of the BRAF gene in human cancer. Nature 417, 949–954. 10.1038/nature00766.

79. O’Shea, T.M., Ao, Y., Wang, S., Ren, Y., Cheng, A.M., et al. (2024). Derivation and transcriptional reprogramming of border-forming wound repair astrocytes after spinal cord injury or stroke in mice. Nature Neuroscience.

80. Squair, J.W., Milano, M., de Coucy, A., Gautier, M., Skinnider, M.A., et al. (2023). Recovery of walking after paralysis by regenerating characterized neurons to their natural target region. Science 381, 1338–1345. 10.1126/science.adi6412.

